# Genome sequencing of 196 *Treponema pallidum* strains from six continents reveals additional variability in vaccine candidate genes and dominance of Nichols clade strains in Madagascar

**DOI:** 10.1101/2021.08.17.456619

**Authors:** Nicole A.P. Lieberman, Michelle J. Lin, Hong Xie, Lasata Shrestha, Tien Nguyen, Meei-Li Huang, Austin M. Haynes, Emily Romeis, Qian-Qiu Wang, Rui-Li Zhang, Cai-Xia Kou, Giulia Ciccarese, Ivano Dal Conte, Marco Cusini, Francesco Drago, Shu-ichi Nakayama, Kenichi Lee, Makoto Ohnishi, Kelika A. Konda, Silver K. Vargas, Maria Eguiluz, Carlos F. Caceres, Jeffrey D. Klausner, Oriol Mitjà, Anne Rompalo, Fiona Mulcahy, Edward W. Hook, Sheila A. Lukehart, Amanda M. Casto, Pavitra Roychoudhury, Frank DiMaio, Lorenzo Giacani, Alexander L. Greninger

## Abstract

In spite of its immutable susceptibility to penicillin, *Treponema pallidum* (*T. pallidum*) subsp. *pallidum* continues to cause millions of cases of syphilis each year worldwide, resulting in significant morbidity and mortality and underscoring the urgency of developing an effective vaccine to curtail the spread of the infection. Several technical challenges, including absence of an *in vitro* culture system until very recently, have hampered efforts to catalog the diversity of strains collected worldwide. Here, we provide near-complete genomes from 196 *T. pallidum* strains – including 191 *T. pallidum* subsp. *pallidum* – sequenced directly from patient samples collected from 8 countries and 6 continents. Maximum likelihood phylogeny revealed that samples from most sites were predominantly SS14 clade. However, 99% (84/85) of the samples from Madagascar formed two of the five distinct Nichols subclades. Although recombination was uncommon in the evolution of modern circulating strains, we found multiple putative recombination events between *T. pallidum* subsp. *pallidum* and subsp. *endemicum*, shaping the genomes of several subclades. Temporal analysis dated the most recent common ancestor of Nichols and SS14 clades to 1717 (95% HPD: 1543-1869), in agreement with other recent studies. Rates of SNP accumulation varied significantly among subclades, particularly among different Nichols subclades, and was associated in the Nichols A subclade with a C394F substitution in TP0380, a ERCC3-like DNA repair helicase. Our data highlight the role played by variation in genes encoding putative surface-exposed outer membrane proteins in defining separate lineages, and provide a critical resource for the design of broadly protective syphilis vaccines targeting surface antigens.

**Author Summary:** Each year, millions of new cases of venereal and congenital syphilis, caused by the bacterium *Treponema pallidum* (*T. pallidum*) subsp. *pallidum,* are diagnosed worldwide, resulting in significant morbidity and mortality. Alongside endemic circulation of syphilis in low-income countries, disease resurgence in high-income nations has underscored the need for a vaccine. Due to prior technological limitations in culturing and sequencing the organism, the extent of the genetic diversity within modern strains of *T. pallidum* subsp. *pallidum* remains poorly understood, hampering development of a broadly protective vaccine. In this study, we obtained 196 near-complete *T. pallidum* genomes directly from clinical swabs from eight countries across six continents. Of these, 191 were identified as *T. pallidum* subsp. *pallidum*, including 90 Nichols clade genomes. Bayesian analysis revealed a high degree of variance in mutation rate among subclades. Interestingly, a Nichols subclade with a particularly high mutation rate harbors a non-synonymous mutation in a putative DNA repair helicase. Coupling sequencing data with protein structure prediction, we identified multiple novel amino acid variants in several proteins previously identified as potential vaccine candidates. Our data help inform current efforts to develop a broadly protective syphilis vaccine.

## Introduction

Syphilis, caused by the spirochete bacterium *Treponema pallidum* subspecies *pallidum* (TPA) remains endemic in low-income countries, where the majority of cases of this infection occurs. A surge in syphilis incidence, however, has been recorded as well in mid- and high-income nations, primarily among men who have sex with men (MSM) and persons living with HIV (PLHIV). The United States saw a 6.5-fold increase in primary and secondary syphilis cases between 2000 and 2019 (1,2), driven in large part by cases among MSM, although cases among heterosexual individuals are now rising rapidly as well. Globally, there were approximately 6 million new cases per year among 15-49 year olds in 2016 (3). One million of these cases occur in pregnant women, of which 63% are in sub-Saharan Africa alone (4). Preventing cases among women of childbearing age is a critical worldwide public health initiative, as TPA can cross the placenta and cause spontaneous abortion and stillbirth. Maternal-fetal transmission of syphilis caused approximately 661,000 adverse birth outcomes globally in 2016 alone (5). In the United States, congenital syphilis is also rising, from 9.2 per 100,000 live births in 2013 to 48.5 per 100,000 live births in 2019, more than a five-fold increase (2).

Given rising infection rates, increasing difficulties in procuring benzathine penicillin G (BPG) for treatment (6), and widespread *T. pallidum* resistance to azithromycin (7–9) which is no longer a viable alternative to BPG, the development of a vaccine against syphilis has become a public health priority. To this end, the syphilis spirochete poses a particular challenge. In contrast to other gram-negative bacteria, TPA has a remarkably low surface density of integral outer membrane proteins (OMPs) (10,11) and uses phase variation (random ON-OFF switching of expression) to further vary its overall surface antigenic profile (12,13). In parallel, this pathogen has evolved a highly efficient gene conversion-based system able to generate millions of variants of the putative surface-exposed loops of the TprK OMP, thus creating an ever-changing target for the host defenses, which fosters immune evasion, pathogen persistence, and re-infection (14–16). Furthermore, TPA cannot be cultured in axenic culture, instead requiring propagation in rabbit testes or, more recently, in co-culture with rabbit epithelial cells (17). This has further hampered efforts to sequence clinical specimens to catalog regions of conservation and diversity, particularly of the OMPs, which is critical for development of an effective vaccine. As of this writing, consensus sequences of only 67 TPA strains have been deposited in INSDC databases, and of these not more than 53 were recovered directly (or following low passage rabbit culture) from clinical specimens. Additional data exist within the Sequence Read Archive (SRA) for up to 600-800 samples annotated as TPA but, without extensive manual curation and reliable assembly pipelines, high-quality data contained within the SRA remain inaccessible to most users. To inform vaccine development efforts, we generated high quality (<1% ambiguous or missing data) near-complete genomes from 196 *T. pallidum* genomes using hybrid capture, enabling direct determination of sequences from clinical specimens without the need for enrichment by culture in rabbits or *in vitro*. These newly available genomes were analyzed to unveil diversity in potential TPA vaccine targets in combination with *in silico* protein folding technology. Our work broadens our understanding of the molecular underpinnings of TPA, and serve as a resource for developing a broadly protective vaccine effective against syphilis.

## Results

### Nichols clade strains are predominant in Madagascar

As part of ongoing efforts to catalog global *T. pallidum* genomic diversity, we received samples containing *T. pallidum* genomic DNA recovered from primary or secondary lesions. We attempted whole genome sequencing on those with > 100 copies of *tp0574* per 17.5 μL genomic DNA and obtained 196 high quality genomes consisting of <1% ambiguities using a custom hybridization capture panel to enrich for *T. pallidum* DNA followed by processing through a custom bioinformatic pipeline for consensus genome calling involving both de novo assembly and reference mapping to the SS14 reference genome (NC_021508.1). Summary demographic characteristics for all samples, including 191 TPA, four *T. pallidum* subsp. *endemicum* (TEN), and one *T. pallidum* subsp. *pertenue* (TPE) used in this study are presented in Table 1. Samples were collected between 1998 and 2020 from 8 countries (Peru, Ireland, USA, Papua New Guinea, Madagascar, Italy, Japan, and China) across 6 continents. Median coverage of the reference genome by trimmed, deduplicated reads was 76.8x (range 16.5 - 1293.4), with a minimum of 6 reads required to unambiguously call a base. Median input genomes, as determined by *tp0574* qRT-PCR, for successful genome recovery was 4,319 copies (range 101 – 304,484) (Supporting Information 1 – Sample Statistics). The median number of ambiguities in the finished genomes prior to masking was 49 (range 0-10,753).

**Table 1:** Demographic information of samples sequenced in this study.

| Country | Year(s) of Collection | Number of Samples | Sex (n) |  |  | Stage (n) |  |  | Previous Study |
| --- | --- | --- | --- | --- | --- | --- | --- | --- | --- |
|  |  |  | Male | Female | Unknown | Primary | Secondary | Unknown |  |
| Peru | 2019 | 9 | 5 | 0 | 4 | 5 | 0 | 4 | n/a |
| Ireland | 2002 | 11 | 0 | 0 | 11 | 0 | 0 | 11 | (18–20) |
| USA | 1998-2002 | 15 | 11 | 3 | 1 | 14 | 1 | 0 | (19,20) |
| Papua New Guinea | 2019 | 1 | 0 | 0 | 1 | 0 | 0 | 1 | n/a |
| Madagascar | 2000-2007 | 85 | 0 | 0 | 85 | 10 | 16 | 59 | (20–22) |
| Italy | 2017 | 10 | 10 | 0 | 0 | 10 | 0 | 0 | n/a |
| Japan | 2019-2020 | 57 | 34 | 22 | 1 | 0 | 0 | 57 | n/a |
| China | 2018 | 8 | 0 | 0 | 8 | 0 | 0 | 8 | n/a |
| **TOTAL** | **1998-2020** | **196** | **60** | **25** | **111** | **39** | **17** | **140** |  |

Following the assembly of the 196 genomes, we combined our strains with an additional 55 publicly available consensus genomes, including five TPE, two TEN, 11 TPA laboratory isolates highly passaged in rabbit, and 37 direct clinical specimens/low passage rabbit TPA strains. Due to differences in library preparation expected to affect performance of our assembly pipeline, we chose not to reassemble genomes that did not have an available consensus sequence. All genomes were masked at the intra-rRNA tRNA-Ala and tRNA-Ile and highly repetitive *arp* and *tp0470* genes for which short read Illumina sequencing could not resolve position or relative length. Genomes were further masked at all paralogous *tpr* genes prior to recombination masking by Gubbins (23).

The maximum-likelihood phylogenetic tree shown in Figure 1 (and in tabular format in Supporting Information 1 – Sample Metadata) is defined by approximately 130-150 non-recombining SNPs separating any two Nichols and SS14 tips and approximately 1,200-1,450 SNPs separating any two TPA and TPE or TEN tips. It recapitulates several features seen in previous phylogenies of *T. pallidum*. Notably, it includes a SS14 Omega node that contains nearly all SS14 clade samples, as well as tight geographic clustering of samples predominantly collected in China and Japan (24) and characterized by uniform azithromycin resistance caused by the A2058G mutation in the 23S rRNA allele (Figure 1A-C; SS14 Omega – East Asia node). None of these samples was resistant via the A2059G allele. We also observed genotypic azithromycin resistance in geographically diverse samples in both the SS14 and Nichols clades, further supporting the hypothesis that this mutation arises spontaneously (24).

**Figure 1:**
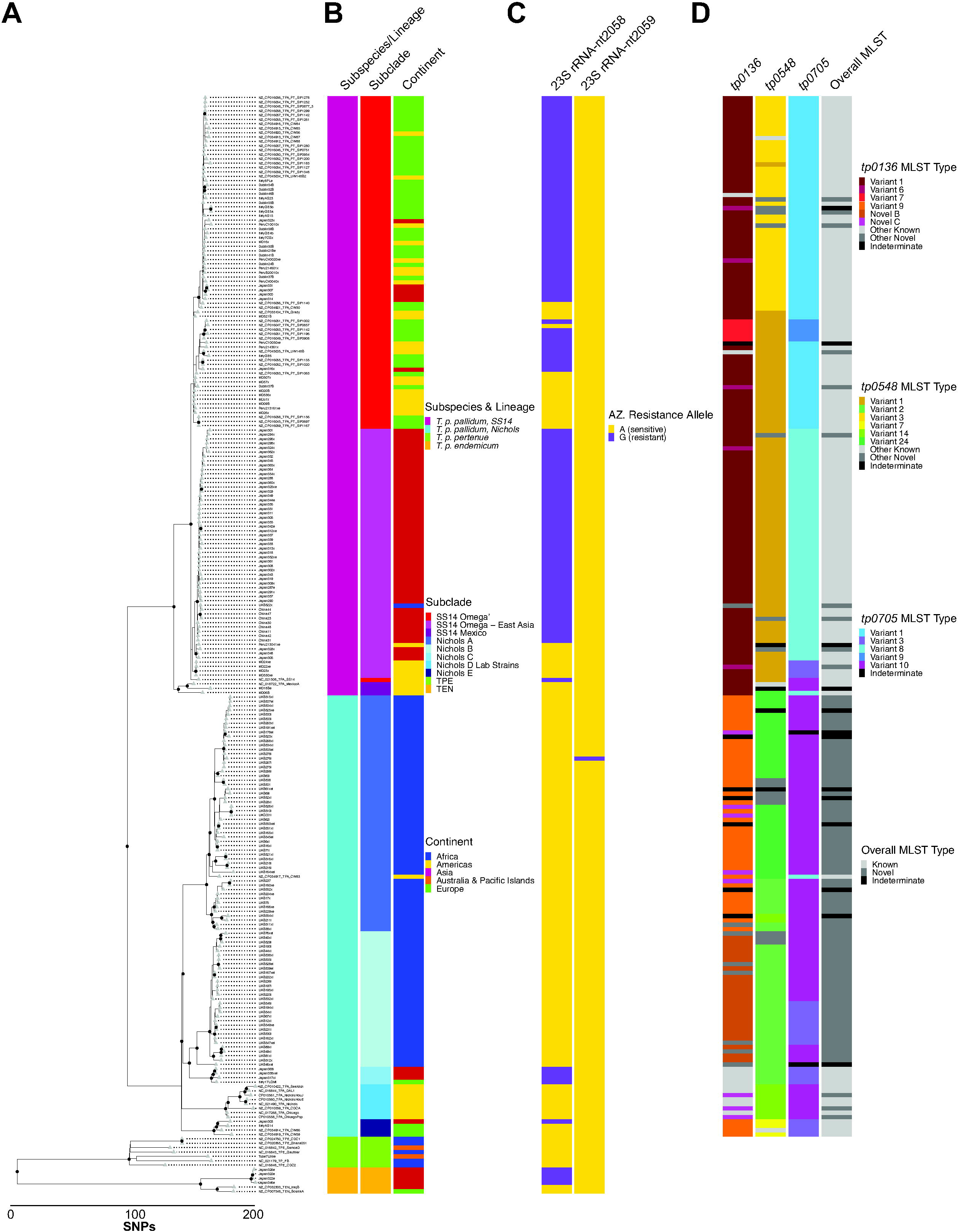
Whole genome phylogeny of *T. pallidum* patient isolates. A) Whole genomes were MAFFT-aligned, recombination-masked, and maximum-likelihood phylogeny determined. Tips are shown as grey triangles and nodes with >0.95 support from 1000 ultrafast bootstraps shown as black circles. B) Subspecies/lineage, subclade, and continent of origin of all samples included in phylogeny. C) Azithromycin sensitivity/resistance as conferred by the 23S rRNA 2058/2059 alleles. Data represents alleles at both rRNA loci. D) MLST subtypes, including novel sequences, for *tp0136*, *tp0548*, and *tp0705*, as well as whether the three alleles constitute a known or novel MLST. Top 6 most abundant sequences at each locus are colored, while other less abundant known and novel sequences are grouped and colored in light and medium grey, respectively. Sequences containing N bases are denoted as indeterminate and shown in dark grey. Expanded metadata for all samples is included in Supporting Information 1.

The most striking feature of our *T. pallidum* phylogeny is the extensive circulation of strains belonging to the Nichols clade in Madagascar. All but one of the 85 Madagascar strains belonged to one of two Nichols subclades, A and B. The former consists of only Madagascar strains except for a single strain from Cuba, and the latter containing only Madagascar samples. Except for an A2059G 23S rRNA mutation (25) observed in one sample, all Madagascar Nichols strains were azithromycin sensitive.

In addition to Nichols subclades A and B primarily from Madagascar, three additional distinct subclades were observed, containing samples collected throughout the world. The Nichols C subclade shares a common ancestor with the Nichols B subclade and is uniformly azithromycin resistant, in contrast to most other Nichols clade samples. Both Nichols D (which contains all laboratory strains) and Nichols E subclades are more distantly related to the Madagascar samples in Nichols A and B. The Nichols E subclade includes two previously reported samples from France as well as two newly sequenced samples from Japan and Italy. Interestingly, both the Japanese and Italian patients whose samples are included in this subclade report their sexual orientation as MSM; one French sample, CW59, was collected from an anal smear. Although this is hardly conclusive, the appearance of distinct TPA clades circulating among MSM individuals has recently been documented in Japan (26), suggesting that this phenomenon may be occurring worldwide.

In addition to the unexpected number of Nichols clade samples, we were also surprised to observe two samples from Maryland that clustered with the very distant SS14 clade genome, MexicoA, originally collected in 1953 from a male living in Mexico. These strains, MD06 and MD18B, diverge from the MexicoA strain by 33 and 24 non-recombining SNPs, respectively; from SS14 Omega strains by about 45 SNPs; and Nichols clade strains by about 150 SNPs. The MexicoA strain is unique in that it shares signatures of both syphilis and yaws organisms in several virulence factors (27,28). To our knowledge, clinical specimens clustering with the MexicoA strain have not previously been reported. Although the MD18B and MD06 samples were collected in 1998 and 2002, respectively, and little demographic information for the samples exists, this is further evidence that our definitions of subspecies of *T. pallidum* may need periodic revisiting.

We also found that four Japanese samples that were clinically diagnosed as syphilis but, based on our whole genome analysis, appear to be part of the TEN subspecies, sharing an ancestor with the canonical TEN genomes IraqB and BosniaA. The observation of TEN samples following a syphilis diagnosis has been previously reported in Japan (29), Cuba (30), and France (presumed to have been contracted in Pakistan) (31). While both the previously discovered and new Japanese TEN samples are resistant to azithromycin via the canonical mutation in the 23S rRNA alleles, neither the Cuban nor French samples are resistant. Furthermore, three of the four samples reported herein were collected from individuals with diverse travel histories (China, Japan, and the Philippines), suggesting that a sexually transmitted TEN outbreak may be even more widespread than previously suspected.

As a mechanism to begin cataloging diversity of several genes known to be polyallelic, we also examined the multi-locus sequence type (MLST) types (32) of all TPA samples using whole genome sequence as well as the combined MLST. Figure 1D highlights the six most common MLST at each locus for TPA samples and whether the overall subtype had been previously reported. Complete MLST data can be found in Supporting Information 1. Across all 238 TPA samples, we found a total of 15 unique complete *tp0136* sequences, including six not previously reported in the MLST database, which contains 26 alleles. Seventeen unique *tp0548* sequences were found, including seven novel sequences, relative to 58 known alleles. All five observed *tp0705* alleles had been previously reported. In total, excluding the 13 samples that were indeterminate at any of the three loci, we found a total of 40 unique haplotypes, including 22 not previously reported in the MLST database. Overall, 88 of 225 *T. pallidum* subsp. *pallidum* samples had a novel overall haplotype, including at least one sample from each country from which samples were obtained, and all 76 (100%) from Madagascar, underscoring the importance of wide geographic sampling to catalog the diversity of TPA strains.

### Putative recombination shapes modern *T. pallidum* subsp. *pallidum* genomes but remains a rare event

Although *T. pallidum* does not have any known plasmids or infecting phages, recombination has nonetheless been shown to be an important mechanism by which genetic diversity may be generated in this pathogen (27,33,34). In particular, the *tpr* family of paralogs is thought to have arisen through gene duplication (28,35); for this reason, all *tpr* genes have been masked for all analyses in this study. Figure 2A shows the comparison of ML tree topology between genomes that have been recombination masked (left) or unmasked (right) (with tip order included in Supporting Information 2). Although no samples were classified to different subclades when recombinant loci were not masked, the overall tree topology was altered. Notably, the SS14 Omega node had more distinct subclades in the absence of recombination masking, suggesting that much of the diversity within SS14 Omega is due to recombination rather than mutation. Furthermore, the Nichols B subclade of Madagascar samples becomes the outgroup within the Nichols clade in the absence of recombination masking.

**Figure 2:**
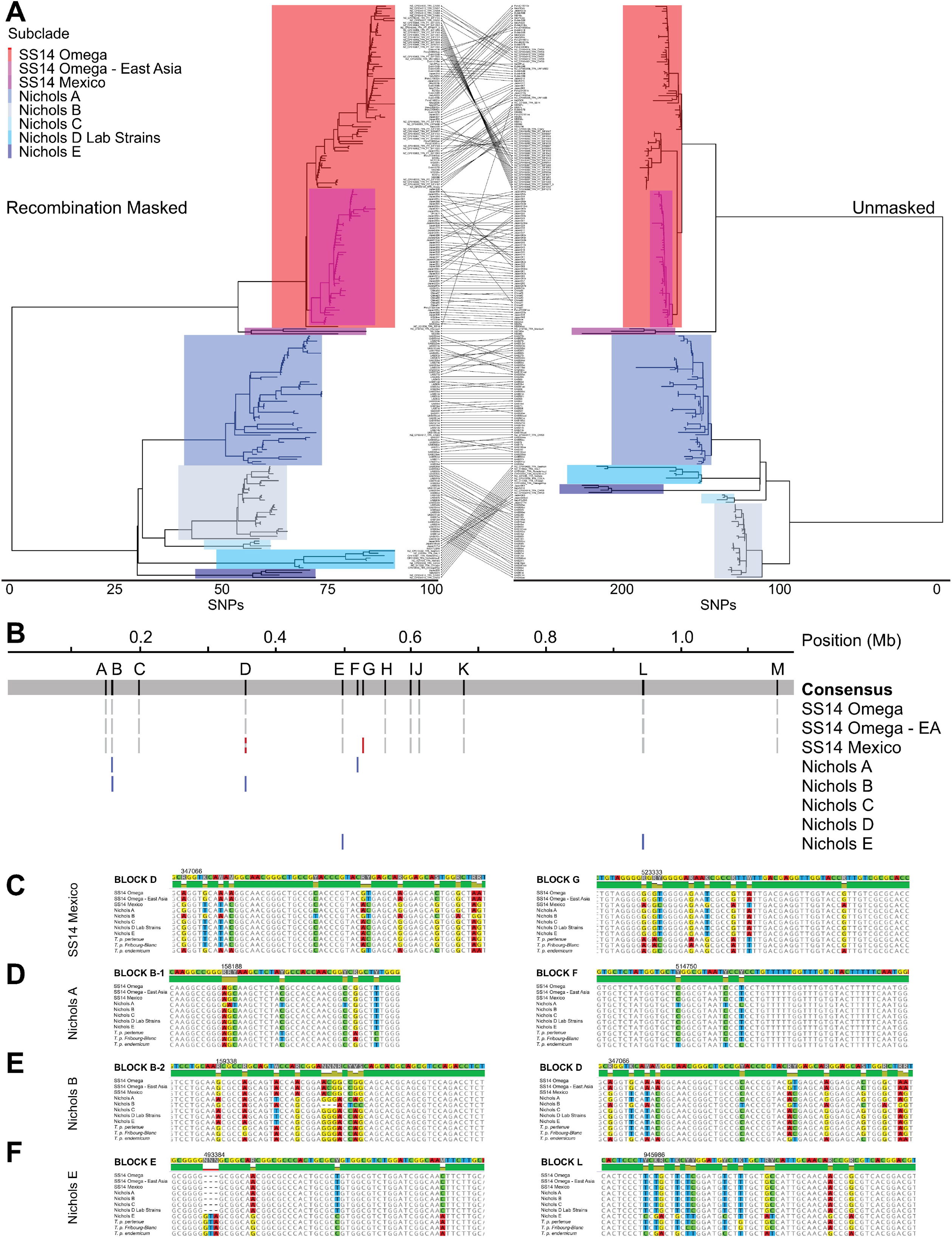
Effect of recombination on *T. pallidum* subsp. *pallidum* evolution. A) Recombination-masked (left) and unmasked (right) phylogenies, with equivalent subclades highlighted. Relative position of each tip is traced between the two panels. B) Putative recombinogenic regions in each clade. Genomic position is relative to the length of the MAFFT alignment. Consensus alignment of all tips is shown on the grey panel, with recombination blocks lettered above. Grey blocks represent recombination that occurred during evolution of the SS14 clade. Red and blue blocks represent recombination events unique to each clade. Mixed grey and colored blocks are regions of ancestral recombination that had a second event unique to that clade. C-F) Two example regions of recombination in SS14 Mexico (C), Nichols A (D), Nichols B (E), and Nichols E (F). Genomic position of the first divergent base in the window shown are shown with NC_021508.1 numbering.

The method of recombination detection we employed relies on identification of an increased density of SNPs per sliding window throughout the clonal frame rather than identification of a discrete donor for each putative recombination event. Although previous analyses have found more recombination in the Nichols clade than SS14 (33), our use of more than 90 clinical specimens belonging to multiple Nichols subclades, albeit with a geographic bias, likely provides a more complete picture of the evolutionary processes that shaped the Nichols clade. In spite of the number of samples examined, recombination remained a rare event in Tpr-masked genomes. Of the 474 nodes on the ML tree, including 238 tips and 236 internal nodes, only 27 branches with recombination were detected. Sixteen of these were on internal nodes and 11 on extant. Of the extant recombination events, four were detected in the 101 Nichols clade samples, and seven of 137 in the SS14 clade samples, suggesting no clade-specific differences in recombination (*p*=0.7633, Fisher’s Exact test).

Figure 2B highlights the positions of identified recombinant regions in the aligned genomes, with grey blocks corresponding to recombination that occurred during the separation of SS14 and Nichols clades, and colored blocks corresponding to recombination events that occurred during the evolution of individual subclades. The grey and red striped block represents a second recombination event in the SS14 Mexico clade that occurred in the same region as the ancestral recombination. As has been previously reported (33,36), many of the identified recombinant regions correspond to the most diverse genes in *T. pallidum*, such as *tp0136*, *tp0326 (BamA)*, and *tp0515 (LptD)* (Supporting Information 2). Notably, many of the ORFs identified as recombinant encode proteins that are predicted to be at least partially surface exposed, and therefore the increased SNP density may represent either bona fide recombination or selective pressure of the host immune system on non-recombinant genes.

Recombination events specific to each subclade shown in Figure 2B were examined, with representative data in Figure 2C-F for SS14 Mexico, Nichols A, Nichols B, and Nichols E subclades, respectively. Windows of approximately 60 bases of the alignments of putative recombinant regions are shown, and include additional *T. pallidum* species members TPE, TEN, and the *T. pallidum* Fribourg-Blanc treponeme, recently proposed to be reclassified as a TPE strain, due to its genetic similarity to other yaws strains (37), with non-identical nucleotides highlighted. Interestingly, several of the identified recombinant loci, including Block G in SS14 Mexico, Block F in Nichols A, and Blocks E and L in Nichols E, have sequences identical to those found in all 6 TEN genomes included in Figure 1. TEN or TPE sequences have been found previously in several Nichols clade samples, suggesting prior recombination (33,36). However, our markedly extended phylogeny of the Nichols clade suggests that recombination between TPA and TEN has independently occurred on multiple occasions. This demonstrates that inter-subspecies recombination continues to play an important role in the diversification of *T. pallidum* subspecies.

### *T. pallidum* subsp. *pallidum* subclades have different rates of SNP accumulation

The evolutionary history of TPA has been a point of considerable debate in recent years, particularly in light of new evidence that could not exclude the presence of TPA in Northern Europe in the late 15^th^ century, casting doubt on the popular theory that venereal syphilis was introduced to Europe by the returning Columbian expeditions (36). In order to determine the date of the most recent common ancestor (MRCA) of the samples included in our study, we first analyzed the temporal signal present among TPA strains by regressing the root-to-tip distances in the SNP-only maximum-likelihood tree (Figure 3A). The left panel shows this calculation performed on a tree that included 11 highly passaged laboratory strains (eight in Nichols clade and three in SS14 clade), identified by open circles, while the right panel is based on a tree that excluded laboratory strains. Notably, the negative slope seen for the Nichols clade appears to be due to the presence of laboratory strains. This is consistent with accelerated accumulation of SNPs during routine passage of the laboratory strains for decades between collection and sequencing. Therefore, laboratory strains were excluded from further dating analysis.

**Figure 3:**
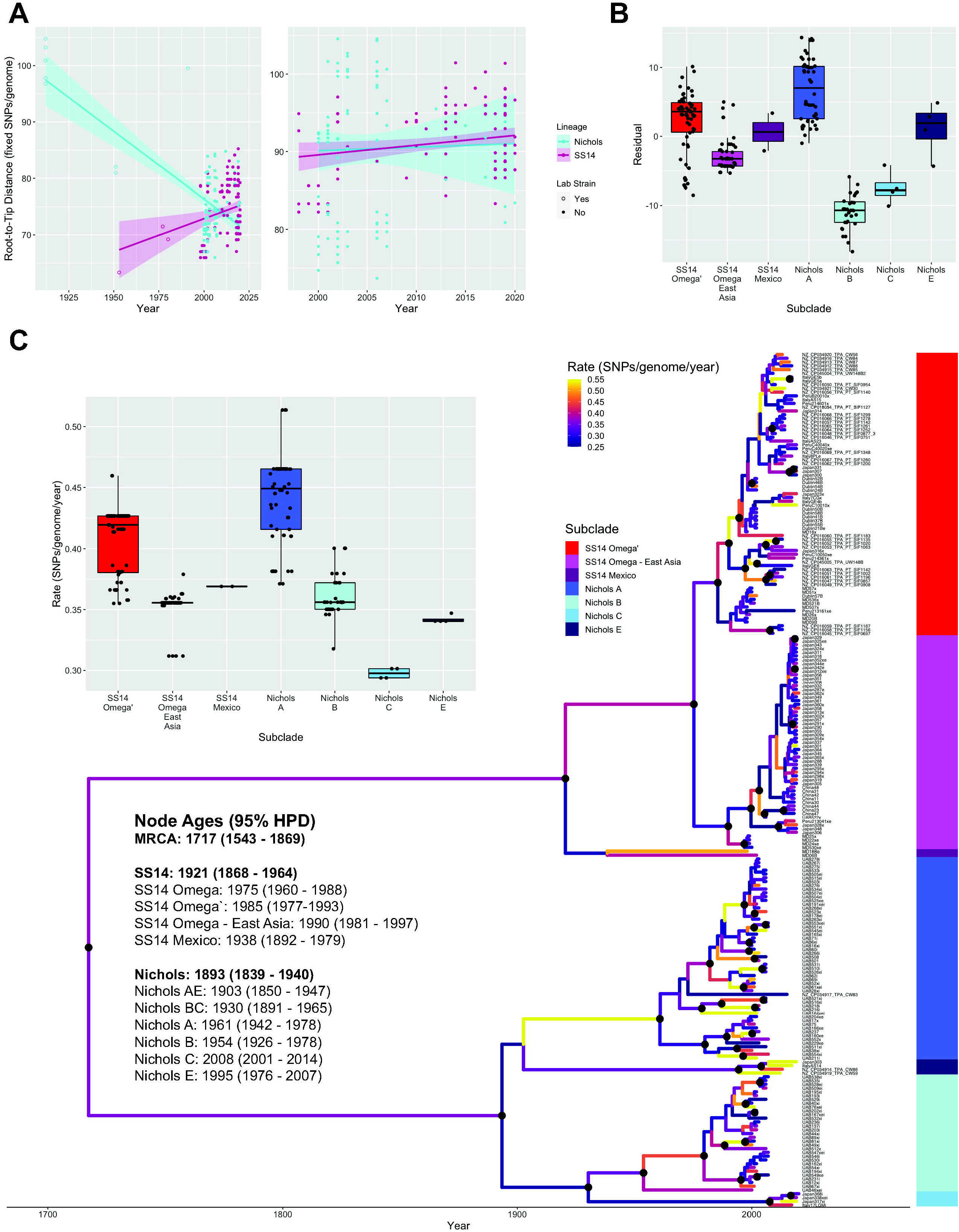
SS14 and Nichols subclades have different rates of SNP accumulation. A) Linear regressions for recombination-masked root-to-tip distances from maximum likelihood phylogeny as a function of year of collection, including (left) or not including (right) highly passaged laboratory strains. B) Residuals from linear regression without laboratory strains were plotted per subclade, *p* < 2e^-16^, ANOVA. C) Bayesian maximum clade credibility tree showing mean common ancestor heights. Highlighted nodes have a posterior probability of >0.95, and branch colors reveal rate of change (SNPs per genome per year). Ages and 95% highest posterior density are included for nodes of interest including the TPA, SS14, and Nichols ancestral nodes, as well as those of each subclade. Inset: For each tip, mean rates of SNP accumulation along branches with >0.95 posterior probability were plotted per subclade, *p* < 2e^-16^, ANOVA.

We were curious as to why the Pearson correlation coefficients of the SS14 and Nichols clades (0.200 and 0.023, respectively) were so poor even in the absence of laboratory strains, and hypothesized that this may be due to differences inherent to the polyphyletic structure of both clades. We tested this by plotting the residuals of the regression by subclade and found significant differences between groups (Figure 3B, *p* < 2e^-16^, ANOVA), suggesting that rates of SNP accumulation may differ across the TPA phylogeny.

Therefore, we proceeded to Bayesian ancestral reconstruction and dating of clinical specimens by BEAST 2 (38), using an uncorrelated relaxed clock with a starting rate of 3.6×10^-4^ (24,39) as a prior model to account for differences in rates of mutation in different branches of the tree. Figure 3C shows the dated Bayesian phylogeny, with branches colored to reflect the rate of SNP accumulation. Black nodes have a posterior probability of >95%. Consistent with previous studies (24,36,39), we dated the MRCA of TPA to 1717 (95% HPD 1543-1869), the Nichols clade to 1893 (1839-1940), and the SS14 clade to 1921 (1868-1964), and found that the rates of SNP accumulation on branches with >95% posterior probability ranged between 0.2 and 0.73 fixed SNPs/year. The inset figure shows the mean rates of diversification on branches with >95% posterior support for each tip, supporting our hypothesis that different subclades have different rates of mutation (*p* < 2e^-16^, ANOVA).

### Host immune pressure drives mutation in the same putative antigens in SS14 and Nichols clades

Observed differences in accumulation of SNPs among subclades may represent the effects of sampling bias or bottlenecks or may reflect differences in the underlying biology. To examine the functional differences that define each subclade (including loci identified as recombinant (Figure 2), we used augur (40) to reconstruct the ancestral nodes identified in the recombination-masked ML phylogeny, transferred ORF annotations from the TPA reference genome NC_021508.1, and translated each ORF to detect coding changes. Figure 4A shows all nodes used for these analyses, with subclade tips collapsed for simplicity. All coding changes detected in Node 101 (SS14 clade ancestral) relative to Node 001 (Nichols clade ancestral, considered equivalent to the TPA root node for these calculations) are shown; data for all additional parent-child node pairs are included in Supporting Information 3. Forty-nine of 1002 putative ORFs were altered between SS14 and Nichols ancestral nodes, with a total of 134 non-synonymous mutation events. We defined a mutation event as a single amino acid change, insertion/deletion, or frameshift. We did not separately include the effects of putative recombination events because we did not attempt to formally characterize recombination donors, and therefore could not disentangle the effects of recombination from selective pressure driving increased mutation.

**Figure 4:**
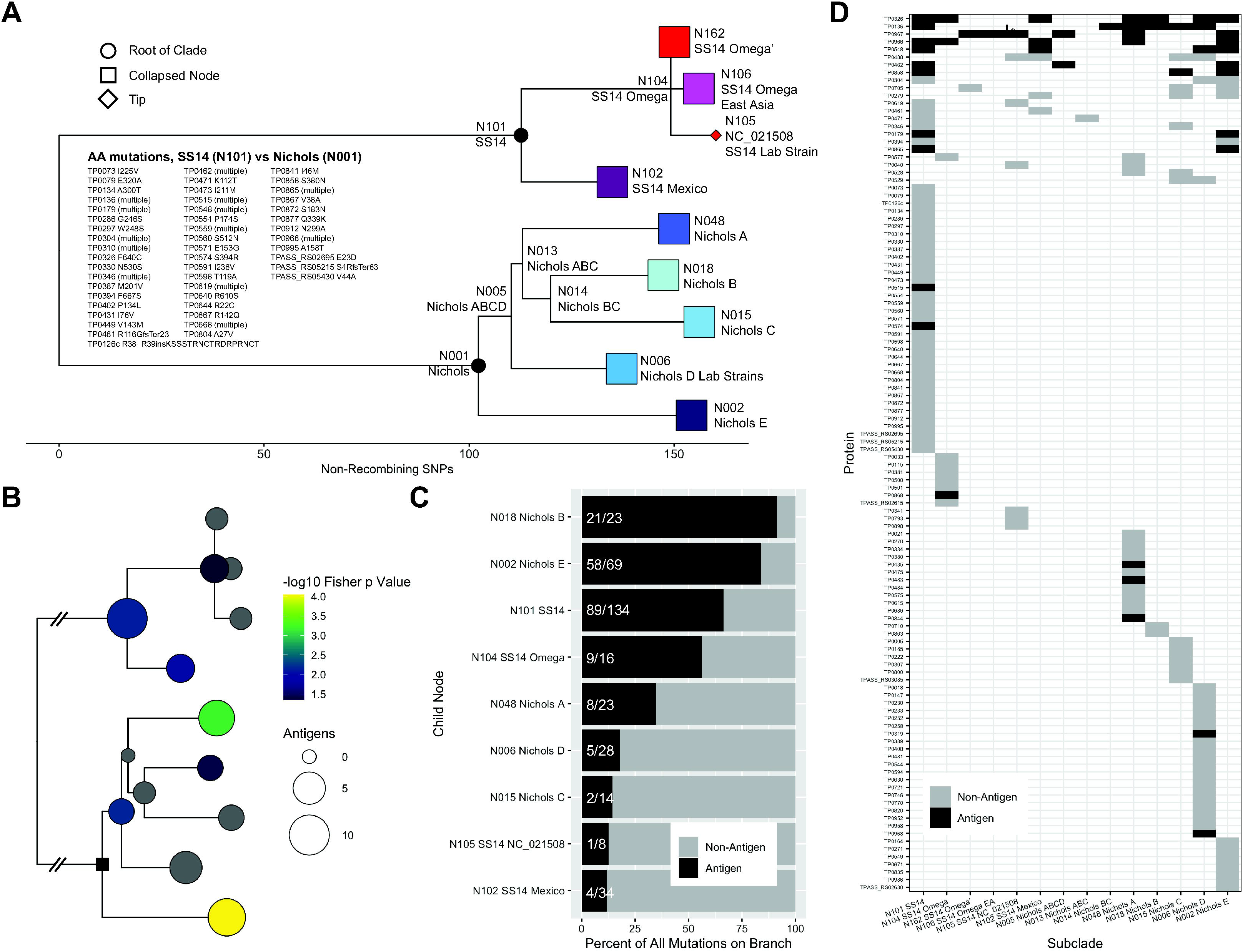
Coding mutations in the *T. pallidum* subsp. *pallidum* phylogeny. A) Whole genome ML phylogeny of TPA, with tips collapsed to the subclade node. Open reading frames of inferred ancestral sequences for each node were annotated based on the SS14 reference sequence NC_021508. Coding mutations, including for putative recombinant genes, for each child node were determined relative to its parent node (complete list in Supporting Information 4). Loci with amino acid differences (n=49 loci, n=134 individual AA mutation events) in the SS14 ancestral clade node (N101) are shown relative to the Nichols ancestral node (N001). B) Positions are equivalent to those shown in A. Black square represents the Nichols Ancestral Node (N001). Number of antigens with coding mutations on each child node relative to parent node. Color represents p value of for overrepresentation by Fisher’s Exact test of antigens among all mutated proteins per branch; those in grey have a p value > 0.05. C) Percentage of total individual mutation events per branch. Raw numbers of mutation events in antigens per total mutation events are shown for each branch. D) Tile plot showing mutated proteins in the ancestral node for each subclade relative to its parent node, colored by antigen or not. Proteins are arranged by number of subclades bearing mutations. Data is recapitulated in Supporting Information 3

We next attempted to define functional changes between the SS14 and Nichols clades by examining overrepresentation of altered loci in categories annotated by structural similarity (41). We used the annotation of the Protein Data Bank (PDB) structure of the highest scoring model, with a confidence cutoff of 75%, allowing 798 coding sequences (CDSs) to be assigned to a total of 62 unique PDB categories. We then performed Fisher’s exact tests to test for overrepresentation of altered proteins in each category. For SS14 vs Nichols ancestral nodes (101 vs 001, Supplementary Figure 1), we only found significant overrepresentation in a single category, “Signaling Protein”, with 3 (*tp0073, tp0640,* and *tp0995*) loci out of the 16 in the category altered. However, because these annotations are by structural similarity rather than known function, it is likely that testing for overrepresentation of structural annotations does not fully capture the functional differences between any two clades.

Because functional annotation of *T. pallidum* proteins is still hampered by the absence of a reverse genetics system, we chose next to focus on alteration of proteins known or suspected to interact with the host immune system. We included proteins that reacted with pooled sera from individuals with known syphilis infection (42,43) or otherwise known to be surface-exposed (Supporting Information 3 - Antigens) and again performed overrepresentation tests (Figure 4B). Along branches with more than 10 altered proteins, only two nodes (N015, Nichols C, and N005, Nichols D Lab Strains) did not have significant p values (p<0.05) relative to their parent. When examining individual mutation events in nodes with more than four altered proteins, mutation in antigenic proteins represents more than 30% of the amino acid variability in more than half of nodes, and at least 10% in all nodes (Figure 4C). Antigenic proteins are enriched among proteins that become mutated relative to their parent node in multiple subclades, representing separate events (Figure 4D). Furthermore, among antigens that were mutated relative to the parent node in more than one subclade, none was exclusive to either the SS14 or Nichols clade. These data suggest that interaction with the host immune system drives a large proportion of the evolution of both major clades of this pathogen, either via individual SNPs or horizontal gene transfer.

However, although antigens are enriched for non-synonymous mutations relative to the rest of the proteome, mutation of non-antigenic proteins may make considerable contributions to *T. pallidum* pathogenicity and immune interaction. When examining proteins whose mutation was unique to a single clade (Figure 4D), we found a C394F mutation in the ERCC3-like DNA repair helicase TP0380 (44) only in the Nichols A subclade, which had a much higher median rate of SNP accumulation than any other subclade (Figure 3C). It is plausible that mutation of this helicase compromises DNA repair and contributes to a more rapid rate of evolution within this clade.

### Predicted structural changes of putative surface proteins not limited to polymorphic residues

For any protein, multiple independent mutation events along several branches of the TPA phylogeny strongly suggest the protein is under selective pressure. Of the six proteins that undergo mutation along four or more of the 14 branches in the phylogeny (Figure 4D and Supporting Information 3 – Heatmap), five (TP0136, TP0326, TP0548, TP0966, and TP0967) are known to be antigenic. TP0326, TP0548, TP0966, and TP0967 are likely outer membrane proteins based on their homology to *N. gonorrhoeae* BamA (TP0326) and *E. coli* FadL (TP0548) and TolC (TP0966, TP0967) (45) and reviewed in (46). TP0136 is a lipoprotein that appears to be localized to the outer membrane, where it functions as a fibronectin- and laminin-binding adhesin (47–49). To date, recombinant TP0136 and TP0326 have been tested as potential vaccine candidates in rabbits, with TP0136 delaying ulceration but not providing full protection upon challenge (47), and TP0326 providing partial protection in some studies (50,51), while not protective in others (52). Although antigens harboring polymorphisms would not traditionally be considered viable vaccine candidates, the paucity of outer membrane proteins in *T. pallidum* (46) demands evaluation of imperfect candidates.

Accordingly, for the five most frequently mutated putative outer membrane antigens, we developed models that highlight the positions predicted to undergo the most structural change upon mutation, including those at orthogonal sites. We first performed global alignments of sequence variants for each of the five proteins using hhpred (53–55) (Supplementary Figures 2-6, panel A, and Supporting Information 4). We then generated composite homology models of the SS14 variant using RosettaCM (56) guided by hhpred sequence alignment. Ribbon structures and surface contours with highlighted polymorphic residues of the SS14 variant are shown in Supplementary Figures 2-6, panels B and C, respectively. Then, the SS14 model was used as a template for predicting the structure of variants of other strains. We performed a global superposition of variant structures and computed an average per-atom displacement relative to the reference model (taking sequence changes into account, see Methods). The resulting per-atom deviations were then mapped onto the model of the SS14 variant, with blue representing regions of the lowest displacement from the SS14 model and red the highest (Supplementary Figures 2-6, panel D). This approach allowed detection of structural changes not simply at the site of the polymorphism, but also orthogonal changes due to disruption of hydrogen and other bonds. Furthermore, it allows “tuning” of the structural effect of a mutation on each atom, with the mutation of similar residues (such as leucine to isoleucine) resulting in less displacement of each atom than substitution of dissimilar residues (such as arginine to histidine). N terminal residues comprising predicted secretion sequences are not shown for TP0136 (48) or TP0326 (57). Best estimates for Gram negative signal peptides were predicted by SignalP 5.0 (58) for TP0548, TP0966, and TP0967 SS14 variants and excluded from display.

In spite of slightly different approaches employed in their generation and our use of the SS14 variant rather than Nichols, our structural models for TP0326, TP0548, TP0966, and TP0967 generally agree with the models recently proposed by Hawley *et al.* (45). TP0326 is a large multidomain component of the β-barrel Assembly Complex (BAM) and includes a C-terminal β-barrel. Consistent with previous studies (34,45,57,59), we found that extracellular loop (ECL)-4 and the serine-rich tract of ECL-7 contribute to much of the between-strain structural diversity (arrows and single arrowheads, respectively, Supplementary Figure 2B-D). We also found that the large ECL-3 (double arrowheads, Supplementary Figure 2B-D) had nine polymorphic residues, rendering the entire exposed surface of the protein variable due to strain-to-strain variation in ECLs, particularly 3, 4, and 7 (Supplementary Figure 2C-D).

In contrast to TP0326, the structure and function of which has been studied extensively, less is known about TP0548, a predicted homolog of the *E. coli* fatty acid transporter FadL. We predict the structure to be a 14-stranded β-barrel, with periplasmic C-terminal α-helices, consistent with previous studies (45). Prediction of linear B cell epitopes (BCEs) using BepiPred 2.0 (60) revealed that, depending on the as-yet unknown position of the cleavage of the N terminal signal sequence, up to four linear BCEs eight residues or longer are predicted to occur in invariant, extended host-facing loops at the N-terminus and ECL-2 (Supplementary Figure 3A, arrows in Supplementary Figure 3B show the relevant loops), rendering them potentially of use in a vaccine cocktail. Notably, the displacement seen in ECL-2, containing BCEs 3 and 4, (Supplementary Figure 3D, arrow) is most likely due to stochastic differences in predicting the conformation of the flexible loop rather than true structural variation.

Both TP0966 and TP0967 are predicted to be orthologs of the *E. coli* efflux pump TolC (45), and are predicted to have a tri-partite structure, with each monomer contributing four β-strands to β-barrel that spans the outer membrane with BCEs predicted within the ECLs (45). Supplementary Figures 4B and 5B highlight a single monomer for TP0966 and TP0967, respectively; both ECLs of TP0966, and ECL1 of TP0967, contain polymorphic residues that disrupt predicted linear B cell epitopes (Supplementary Figures 4A and 5A, Supplementary Figures 4C and 5C, arrows). For both TP0966 and TP0967, the residues with the most displacement that disrupt the extracellular surface are not the polymorphic positions (Supplementary Figures 4D and 5D, arrows). Rather, in TP0966, the polymorphic charged residues in and adjacent to the ECLs may cause changes to electrostatic interactions that influence loop position. In TP0967, the length of the poly-glycine tract alters the position of ECL1. The likely result is disruption of the conformational epitopes formed by the surface loops in TP0966 and TP0967.

Finally, we generated a structural model of TP0136, and found it to adopt a 7-bladed beta-propeller fold in its N-terminal domain, followed by a relatively unstructured C-terminal domain (Supplementary Figure 6B). The beta-propeller structure is noteworthy as it is homologous to structures found in several eukaryotic integrins that mediate binding to the extracellular matrix (61), as well as to bacterial lectins (62). Several tracts of serine and lysine repeats are a unique structural feature of TP0136; the beta-propeller fold of TP0136 allows these intrinsically disordered regions to form unstructured loops between beta strands. Unsurprisingly, the surfaces that comprise the β-strands have some polymorphisms (Supplementary Figure 6B and C, boxed region) but they are not predicted to cause extensive structural displacement and disruption of the fold, as shown by primarily blue coloring in the boxed region of Supplementary Figure 6D.

Interestingly, the deletion in TP0136 that appears in 4 sequence variants (2, 5, 22, 23, and 24, Supplementary Figure 6A, alignment position 161-192) and entirely removes the large flexible loop annotated by an arrow in Supplementary Figure 6B-D is not found in any ancestral node sequences (Figure 4), but arises independently in strains from multiple geographic locations, including Nichols clade strains from Madagascar and the United States (subclades A, B, and D), and SS14 clade strains from Japan, Peru, and Ireland (subclades Omega – East Asia and SS14 Omegà), consistent with this genomic region being a hotspot for recombination (Figure 2).

## Discussion

In recent years, *T. pallidum* genomics has been significantly advanced by projects aimed at studying the origin and spread of strains responsible for the modern syphilis pandemic (24,26,36,39,63), as well as the emergence of azithromycin resistance (24,26,63). Increasingly the challenge in *T. pallidum* genomics will be attaining complete genomic sequences from undersampled regions, associating genomic sequencing with spirochete biochemical functions, and gaining actionable insights into *T. pallidum* evolution that inform vaccine design.

With these goals, we generated 196 near-complete *T. pallidum* genomes from diverse locations, including three countries – Peru, Italy, and Madagascar – with no previous complete genomes publicly available. Peruvian samples (n=9) belonged exclusively to the SS14 Omegà subclade, which contains samples collected worldwide and corresponds to the largest SS14 sub-lineage in a recent analysis of SNPs in TPA strains (63). Eight of the ten Italian strains also belonged to the Omegà subclade.

The remaining two Italian strains, collected in Turin and Bologna, were of two distinct Nichols subclades, one of which clustered with three Japanese syphilis strains in Nichols subclade C, and the other clustered with samples of Japanese and French origin, forming the distantly related Nichols subclade E. Notably, none of the Japanese or Italian samples clustered with the Malagasy Nichols samples, which, but for a single Cuban strain in Nichols subclade A, formed two private subclades. Because the samples from Madagascar were collected between 2000-2007, it is unknown whether there has been introduction of additional lineages of TPA in the intervening years, or whether the two nearly private subclades are reflective of the currently circulating strains.

Sample collection date is also an important consideration to the interpretation of azithromycin resistance data. None of the strains collected in the USA between 1998 and 2002 were resistant to azithromycin. However, this was prior to the detection of widespread azithromycin resistance in the United States (8); therefore, the lack of resistance detected in the strains sequenced for the present study should not be considered representative of the current status. Only one of the strains collected from Madagascar between 2000 and 2007 was resistant to azithromycin; no subsequent sampling has been performed, thus, no conclusions about azithromycin resistance in strains currently circulating in Madagascar can be drawn.

In our study, as in other recent global *T. pallidum subsp. pallidum* genomics initiatives (24,63), samples were collected and sequenced based on availability rather than representing an even distribution based on global burden of disease. The result of this is that, although we gained a broader picture of worldwide diversity, some regions (North America, western Europe, eastern Asia) continue to be overrepresented, while other regions (Africa – particularly Sub-Saharan Africa, which bears the largest share of cases worldwide – and South Asia and South America) are still vastly under-sampled. However, an important takeaway from our study as well as the recent paper from Beale *et al.* (63) is that the general understanding that SS14 represents the vast majority of circulating strains may require revisiting. Although the island nation of Madagascar is unlikely truly representative of the diversity of strains currently circulating in Sub-Saharan Africa, particularly because the samples are 15-20 years old, our finding that 99% of Malagasy strains belong to the Nichols clade, coupled with Beale *et al*.’s discovery of Nichols strains circulating in Zimbabwe and South Africa (63) strongly suggests widespread circulation of Nichols clade TPA in Africa. Clearly, increased sampling must be a priority to enable understanding of syphilis epidemiology in Africa, and to ensure a vaccine covers strains circulating in the regions most hard hit by the modern pandemic.

Our temporal analysis generally agreed with previous estimates of mutation rate (39,63) in spite of the fact that we used a relaxed, rather than fixed, clock model to determine whether there were differences in the rate of mutation along different branches of the *T. pallidum* subsp. *pallidum* phylogeny, which could indicate either different selection pressures or underlying biological differences contributing to the phenotype. Indeed, we found significant differences in the rates of mutation among the subclades (Figure 3C). The Nichols A subclade was particularly interesting to us, given its high median rate of mutation along branches within the subclade with high posterior support. Notably, when we examined the non-synonymous mutations that defined the Nichols A subclade relative to its ancestral node, shared by Nichols subclades A, B, and C (Figure 4A/D), we found that one of the non-synonymous mutations found only within the Nichols A subclade was in TP0380, a putative ERCC3-like DNA repair helicase that interacts with DNA replication machinery by yeast two-hybrid analysis (41,44,64). Although the functional significance of the C394F mutation (C1181A in *tp0380*) is unknown, it is tempting to speculate that it may directly affect DNA repair. This hypothesis of a potential mutator phenotype in *T. pallidum* can now be examined *in vitro*, given the recent description of the first genetic transformation in *T. pallidum* (65). If TP0380 mutation is indeed responsible for the elevated rate of mutation seen within the Nichols A subclade, the implications for vaccine design may be significant.

By definition, an effective syphilis vaccine needs to protect against most strains circulating where the vaccine is administered. Our work further supports that the majority of non-synonymous mutations that define *T. pallidum* subsp*. pallidum* subclades are in proteins putatively located in the outer membrane, or known to react with serum from syphilis patients (Figure 4) (42,43). These data, along with recent structural modeling of *T. pallidum* outer membrane proteins showing that putative B cell epitopes are primarily found on the protein surface predicted to face the host (45), strongly suggest that immune pressure is the most important driver of mutation in *T. pallidum* subsp. *pallidum*. Indeed, our own structural modeling, which highlights regions with the highest structural displacement due to sequence variability, confirms that the regions with the highest displacement are frequently polyallelic (Supplementary Figures 2-6). Given the paucity of *T. pallidum* outer membrane proteins, and the extensive mutation of predicted epitopes, a multivalent vaccination strategy may engender a polyclonal humoral response capable of neutralizing a wider array of strains, a strategy currently being adopted in our laboratory.

Finally, an important caveat to these data is that, due to their extensive recombination and duplication, we excluded arguably the most important *T. pallidum* proteins that interact with the host immune system, the Tpr family (14,28). Although this approach has been used before to ensure an accurate phylogeny free from the confounding effects of recombination (24,63), as well as to prevent mistakes due to improper resolution of their repetitive elements during de novo assembly (39), an understanding of how the *tpr* genes evolve and influence host immunity is critical to developing an efficacious vaccine to *T. pallidum*. Accordingly, we are currently undertaking additional analyses of the Tpr family in these strains, including the hypervariable regions of TprK.

The data presented in this study represent a step forward toward developing a successful vaccine against syphilis. Alongside increased sequencing of strains from regions without extensive sampling, particularly Africa and South America, improved biophysical and computational methods are necessary to unequivocally determine which proteins are expressed on the surface of the bacterium during human infection. The new system to genetically engineer *T. pallidum* (65) will undoubtedly aid these studies, as well as allow the development of strains to test vaccine candidates in animal experiments. Finally, a successful vaccine must not only be efficacious against all circulating strains, but must also be sufficiently low cost and robust to ambient temperatures to allow distribution in the developing world, which is currently bearing the burden of the modern pandemic.

## Methods

### Ethics Statement

All human samples were collected and deidentified following protocols established at each institution. Samples from Ireland, Madagascar, and USA have been previously published (18–22). IRB protocol numbers for collection of the remaining samples are as follows: China: Nanjing Medical University, 2016□050; Italy: Universities of Turin and Genoa, PR033REG2016, University of Bologna, 2103/2016; Japan: National Institute of Infectious Diseases, 508 and 705; Peru: University of Southern California, HS-21-00353; Papua New Guinea, Lihir Medical Center, Medical Research Advisory Committee of the PNG NDOH No: 17.19. Sequencing of deidentified strains was covered by the University of Washington Institutional Review Board (IRB) protocol number STUDY00000885.

### Library Preparation

Samples were collected and DNA extracted using standard protocols (66). Treponemal burden was assessed by quantitative PCR (qPCR) for *TP47* multiplexed with human β-globin, using primer sequences *TP47*-F: 5’-CAAGTACGAGGGGAACATCGAT, *TP47*-R: 5’: TGATCGCTGACAAGCTTAGG, *TP47*-probe: 5’-6FAM-CGGAGACTCTGATGGATGCTGCAGTT-NFQMGB. Pre-capture libraries were prepared from up to 100 ng input genomic DNA using the Kapa Hyperplus kit (Roche), using a fragmentation time of 8 minutes and standard-chemistry end repair/A-tailing, then ligated to TruSeq adapters (Illumina). Adapter-ligated samples were cleaned with 0.8x Ampure beads (Beckman Coulter) and amplified with barcoded primers for 14-16 cycles, followed by another 0.8x Ampure purification.

### *T. pallidum* capture

Capture of *T. pallidum* genomes was performed according to Integrated DNA Technology’s (IDT’s) xGen Hybridization Capture protocol. Briefly, pools of 3-4 libraries were created by grouping samples with similar treponemal load for a total of 500 ng DNA, and Human Cot 1 DNA and TruSeq blocking oligos (IDT) added prior to vacuum drying. The hybridization master mix, containing biotinylated probes from a custom IDT oPool tiling across the NC_010741.1 reference genome, was then added overnight (>16 hr) at 65C. The following day, streptavidin beads were added to the capture reaction, followed by extensive washing, 14-16 cycles of post-capture amplification, and purification with 0.8× Ampure beads. Pool concentration was determined by Qubit assay (Thermo Fisher) and size verified by Tapestation (Agilent). Libraries were sequenced on a 2×150 paired end run on a HiseqX.

### Fastq processing

Fastqs were processed and genomes assembled using our custom pipeline, available at https://github.com/greninger-lab/Tpallidum_WGS. Paired end reads were adapter- and quality-trimmed by Trimmomatic 0.35 (67), using a 4 base sliding window with average quality of 15 and a minimum length of 20, retaining only paired reads. Trimmed reads were then filtered with bbduk v38.86 (68) in two separate steps. First, reads were filtered very stringently, allowing removal of contaminating non-*T. pallidum* reads, against a reference containing the two rRNA loci, with a 100 bp 5’ and 3’ flank, from each of five reference *T. pallidum* genomes (NC_021508.1 (*T. pallidum* subsp. *pallidum* strain SS14), NC_016842.1 (*T. pallidum* subsp. *pertenue* strain SamoaD), NC_016843.1 (*T. pallidum* subsp. *pertenue* strain Gauthier), NC_021179 (*T. pallidum* strain Fribourg-Blanc treponeme), NZ_CP034918.1 (*T. pallidum* subsp. *pallidum* strain CW65)). We used a kmer size of 31, a Hamming distance of 1, a minimum of 98% of kmers to match reference, and removal of both reads if either does not pass these criteria. Second, unmatched reads from the rRNA filtration step were then filtered against the complete reference genomes that had been masked with N at the rRNA loci, using a kmer size of 31 and a Hamming distance of 2. Matching reads from the two steps were concatenated and used for input for genome mapping and assembly.

### Genome assembly

Filtered reads were mapped to the *T. pallidum* street 14 reference genome, NC_021508.1, using Bowtie2 v2.4.1 (69) with default parameters and coverted to bam with samtools v1.6 (70), followed by deduplication by MarkDuplicates in Picard v2.23.3 (71). Prior to *de novo* assembly, rRNA-stripped reads were filtered with bbduk (68) to remove repetitive regions of the genome, including the repeat regions of the *arp* and *TP0470* genes, as well as *tprC*, *tprD*, and the *tprEGF* and *tprIJ* loci, using a pseudo-kmer size of 45 and Hamming distance of 2. *De novo* assembly was performed using Unicycler v0.4.4 (72) using default settings, with rRNA- and repetitive region-stripped paired fastqs as input. Contigs longer than 200 bp were then mapped back to NC_021508.1 reference genome using bwa-mem 0.7.17-r1188 (73) and a custom R script (74) used to generate a hybrid fasta merging contigs and filling gaps with the reference genome. Deduplicated reads were initially remapped to this hybrid using default Bowtie2 settings, local misalignments corrected with Pilon v1.23.0 (75), and a final Bowtie2 remapping to the Pilon consensus used as input to a custom R script (74) to close gaps and generate a final consensus sequence, with each position called at a threshold of 50% of reads supporting a single base. A minimum of six reads were required to call bases; coverage lower than 6x was left ambiguous by calling “N”. All steps of genome generation were visualized and manually confirmed in Geneious Prime v2020.1.2 (76). Following consensus generation, the tRNA-Ile and tRNA-Ala sites that occur within the rRNA loci were masked to N due to short reads being unable to resolve the order of the sites. Consensus genomes were further masked at the *arp* and *tp0470* repeats and *tprK* variable regions prior to further analysis and deposition in the NCBI genome database.

### Phylogeny

Consensus genomes that had been masked at the *arp* and *tp0470* repeats, intra-rRNA tRNAs, and *tprK* variable regions were further masked to N at all *tpr* genes, which are known to be recombinogenic (35). Masked genomes were aligned with MAFFT v7.271 (77) with a gap open penalty of 2.0 and an offset (gap extension penalty) of 0.123. Aligned genomes were recombination masked using 25 iterations of Gubbins v2.4.1 (23). Recombination masking was performed separately with and without *T. pallidum* subsp. *pertenue* and *T. pallidum* subsp. *endemicum* sequences as appropriate. Recombination masking was mapped back onto whole genome sequences and visualized using maskrc-svg v0.5 (78), and iqtree v2.0.3 (79) used to generate a whole genome maximum likelihood phylogeny using 1000 ultrafast bootstraps and automated selection of the best substitution model. A non-recombination-masked maximum likelihood tree was generated using the same parameters but with the raw MAFFT output. Sequences of *tp0136*, *tp0548*, and *tp0705* were extracted and batch queried using the PubMLST database ((32), accessed 01-22-2021).

### Bayesian Dating

TempEst v1.5.3 (80) was used to calculate root-to-tip distances for the SNP-only maximum likelihood phylogeny calculated with or without *T. pallidum* subsp. *pallidum* laboratory strains, assuming one year uncertainty in strains with collection dates estimated. Regressions of distance vs sample date were performed per clade in R. Bayesian dating was performed in the BEAST2 suite (38) using the recombination masked SNP-only (n=600 sites) alignment of *T. pallidum* subsp. *pallidum*, excluding laboratory strains. Priors included a relaxed clock lognormal model with a starting rate of 3.6×10^-4^ (24,39), constant population size, and a GTR +gamma substitution model. Three separate runs, each with 100,000,000 MCMC cycles were performed and the first 10,000,000 cycles discarded as burn-in. All runs converged and were merged prior to calculation of the maximum clade credibility (MCC) tree.

### Ancestral node reconstruction

Augur v10.1.1 (40) was first used to map all tips without recombination masking onto the whole genome phylogeny generated following recombination masking, ensuring appropriate ancestral relationships unconfounded by recombination, using the “refine” function. The “ancestral” function was next used with default settings to infer ancestral node sequences. Sequences of select nodes were aligned to reference NC_021508.1 using MAFFT as above, and annotations of the reference transferred to ancestral node sequences in Geneious. Pairwise global alignments of protein sequences were performed in R using the Biostrings package (81), and analysis and statistical measurements performed in R using custom scripts.

### Antigens

Antigens were manually curated based on being reactive against *T. pallidum* subsp. *pallidum* positive human sera in either of two previous studies (42,43) or, to control for low expression hampering detection by these in vitro methods, by being selected as likely surface proteins or lipoproteins based on extensive literature searches.

### Structural modeling

Genomes were annotated using Prokka v1.14.6 (82), using the --proteins flag to force annotations to comply with NC_021508.1. Translated coding sequences for vaccine-relevant genes were extracted with a custom R script. Sequences containing ambiguities or truncations likely due to assembly gaps in the genome were manually reviewed in Geneious and excluded from further analysis.

Homology modelling in RosettaCM (56) was used to build initial models of the SS14 variant of each protein. For all sequences collected as part of this study, hhpred (53–55) was used to identify homologous structures, and only those sequences with fulllength alignments (covering >70% of the target) with high probability (>95% hhpred score) were considered for structural modelling. Given these alignments, 100 independent modelling trajectories were carried out for a reference sequence, guided by the top 1-7 templates for each target. We used the following templates in modelling each target: TP0136 used 4a2l and 5oj5; TP0326 used 4k3b and 5d0o; TP0548 used 6h3i; TP0966 used 1yc9, 3d5k, 4k7r, 4mt0, 4mt4, 5azs, and 6u94; TP0967 used 1yc9, 3d5k, 5azp, 5azs, and 6u94. For targets TP0966 and TP0967, modelling was carried out considering the complete homotrimeric configuration, using the symmetry of the templates as a guide.

Following homology modelling, the lowest-energy model was selected and used as a starting point for modelling the mutant sequences. We again used RosettaCM, providing the reference model as the “template” and each mutation as the “target” sequence. For each mutant sequence, three models were predicted and the lowest-energy one was used in analysis of structural deviations.

Structural deviation analysis involved comparing the structures of proteins with different sequences, and standard difference metrics (like backbone RMSd) do not properly report differences in sidechain identities. Instead, we used a “per-atom RMSd” metric, where the structures were first superimposed on the reference structure by aligning common backbone atoms. Then, for each atom in the reference structure, the distance was computed not to a corresponding atom, but rather the closest atom of the same chemical identity (e.g., the oxygens of glutamate and aspartate would map to one another). This was then used to calculate the per-atom and per-residue RMS deviations reported in the manuscript. In this part of the analysis, the homotrimeric configuration of targets TP0966 and TP0967 was again used.

Bacterial signal peptide predictions were performed with SignalP 5.0 (58) Linear B cell epitopes were predicted using the BepiPred-2.0 (60).

### Statistics and Visualization

Unless otherwise noted, all statistical analysis was performed in R v 4.0.0. Phylogenetic trees and metadata were visualized with the R packages ggtree (83), treeio (84), and ggplot (85), multiple sequence alignments by R package ggmsa (86) and figures generated using cowplot and Adobe Illustrator v24.1.3.

## Supporting information

Figure S2BCD

Figure S3BCD

Figure S4BCD

Figure S5BCD

Figure S6BCD

## Data Availability

Paired end reads have been uploaded to the NCBI Sequencing Read Archive, Bioproject PRJNA723099. Consensus genomes have been deposited to NCBI Genome, accession numbers CP073381-CP073576 (Supporting Information 1 - Sample Accessions).

## Acknowledgements

The authors would like to thank the individuals who donated specimens for the studies conducted here.

## Figure Legends

**Supplementary Figure 1: Distribution of putative protein functional annotation based on high-confidence Phyre2 models**. Percent of proteins in each category different between the SS14 ancestral clade node (N101) and the Nichols ancestral node (N001) (orange) were compared to annotations across the whole genome (purple). Overrepresentation was tested by Fisher’s exact test, \**p* < 0.05.

**Supplementary Figure 2: Multiple sequence alignment and structural modeling of TP0326.** A) Multiple sequence alignment for all amino acid sequence variants. Polymorphic residues are highlighted, and positions of extracellular loops 3, 4, and 7 are shown. B) Side (left) and top (right) cartoon representation of TP0326, with a color gradient between blue at the N-terminus to red at the C-terminus. C) Side (left) and top (right) space-filling representation of TP0326, with polymorphic residue positions colored magenta. D) Side (left) and top (right) space-filling representation of TP0326, with atoms colored by average per atom displacement in all variants relative to the SS14 reference sequence. Arrows, single arrowheads, and double arrowheads point to positions of ECLs 4, 7, and 3, respectively.

**Supplementary Figure 3: Multiple sequence alignment and structural modeling of TP0548.** A) Multiple sequence alignment for all amino acid sequence variants. Polymorphic residues are highlighted, and positions of relevant predicted B cell epitopes are shown. B) Side (left) and top (right) cartoon representation of TP0548, with a color gradient between blue at the N-terminus to red at the C-terminus. Arrows point to the flexible loops that contain predicted linear BCEs. C) Side (left) and top (right) space-filling representation of TP0548, with polymorphic residue positions colored magenta. D) Side (left) and top (right) space-filling representation of TP0966, with atoms colored by average per atom displacement in all variants relative to the SS14 reference sequence. Arrow points to ECL-2, which contains two invariant predicted BCEs.

**Supplementary Figure 4: Multiple sequence alignment and structural modeling of TP0966.** A) Multiple sequence alignment for all amino acid sequence variants. Polymorphic residues are highlighted. ECLs 1 and 2 are boxed, and the linear BCEs contained in the SS14 variant (#1) are marked in red. B) Side (left) and top (right) cartoon representation of TP0966, with a color gradient between blue at the N-terminus to red at the C-terminus. C) Side (left) and top (right) space-filling representation of TP0966, with polymorphic residue positions colored magenta. Arrow points to polymorphic residues in surface loops. D) Side (left) and top (right) space-filling representation of TP0966, with atoms colored by average per atom displacement in all variants relative to the SS14 reference sequence. Arrows point to the high displacement, non-polymorphic residues.

**Supplementary Figure 5: Multiple sequence alignment and structural modeling of TP0967.** A) Multiple sequence alignment for all amino acid sequence variants. Polymorphic residues are highlighted. ECL1 is boxed, and the linear BCE contained in the SS14 variant (#4) is marked in red. B) Side (left) and top (right) cartoon representation of TP0967, with a color gradient between blue at the N-terminus to red at the C-terminus. C) Side (left) and top (right) space-filling representation of TP0967, with polymorphic residue positions colored magenta. Arrow points to polymorphic residues in surface loops. D) Side (left) and top (right) space-filling representation of TP0967, with atoms colored by average per atom displacement in all variants relative to the SS14 reference sequence. Arrows point to the high displacement, non-polymorphic residues.

**Supplementary Figure 6: Multiple sequence alignment and structural modeling of TP0136.** A) Multiple sequence alignment for all amino acid sequence variants. Polymorphic residues are highlighted. B) Side (left) and top (right) cartoon representation of TP0136, with a color gradient between blue at the N-terminus to red at the C-terminus. C) Side (left) and top (right) space-filling representation of TP0136, with polymorphic residue positions colored magenta. D) Side (left) and top (right) space-filling representation of TP0136, with atoms colored by average per atom displacement in all variants relative to the SS14 reference sequence. Boxed areas represent regions of low displacement in the β-strands. In all panels, arrows point to the large extracellular loop that is removed in variants found in several subclades.

**Supporting Information 1: Expanded metadata for all samples presented in** Figure 1. ***Sample Statistics:*** “Input Genomes” refers to the number of copies of TP47 (*tp0574*) included in pre-capture library preparation. “Coverage” refers to the average deduplicated read depth at any position in the initial mapping of reads to the reference sequence NC_021508. “Length” is the length of the final consensus genome. “Number Ambiguities” and “Percent Ambiguities” refers to Ns in samples prior to masking. ***Sample Accessions***: NCBI Biosample and assembly accessions per sample. ***Sample Metadata:*** “Tip Number” refers to sample position on the phylogenetic tree in Figure 1, with #1 at the bottom of the page. ***tp0136 MLST, tp0548 MLST, tp0705 MLST:*** *Complete* information on MLST alleles, including how novel sequences relate to the closest known match.

**Supporting Information 2: Expanded Recombination Data from** Figure 2**: *Tip Order:*** Top to bottom order of tips in recombination masked and unmasked phylogenies. ***Recombination Blocks:*** Genes included in each recombination block. ***Blocks per Clade***: Precise locations of recombination events detected per clade.

**Supporting Information 3: Expanded data for** Figure 4**. *Summary Data by Locus:*** Information on number of mutations per locus per branch, including functional annotations. ***Antigens:*** List of loci included as antigens. CDS name and genomic position information is from NC_021508.1. ***Heatmap Data:*** Data included in Figure 4D. “Sum” represents the number of nodes at which there is a change in that locus. ***NXXX vs NYYY:*** Detailed information on all detected mutations per branch, named by parent and child node number.

**Supporting Information 4: Expanded data for Supplementary Figures 2-6.** TP0136, TP0326, TP0548, TP0966, and TP0967 variant amino acid sequence by sample and subclade.

