## Supplementary figures and images for "Genome sequencing of 196 *Treponema pallidum* strains from six continents reveals additional variability in vaccine candidate genes and dominance of Nichols clade strains in Madagascar"

### Figure S1

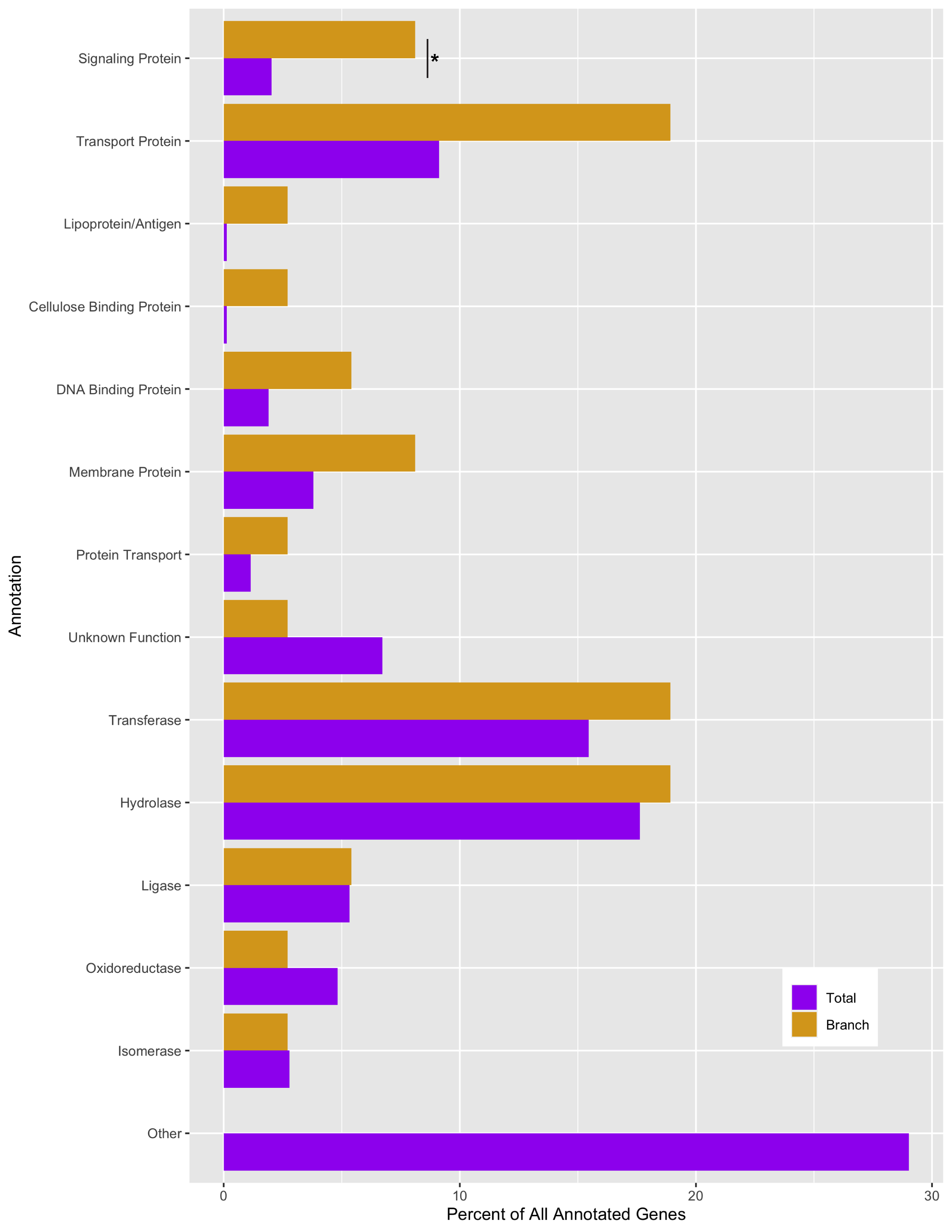

### Figure S2A

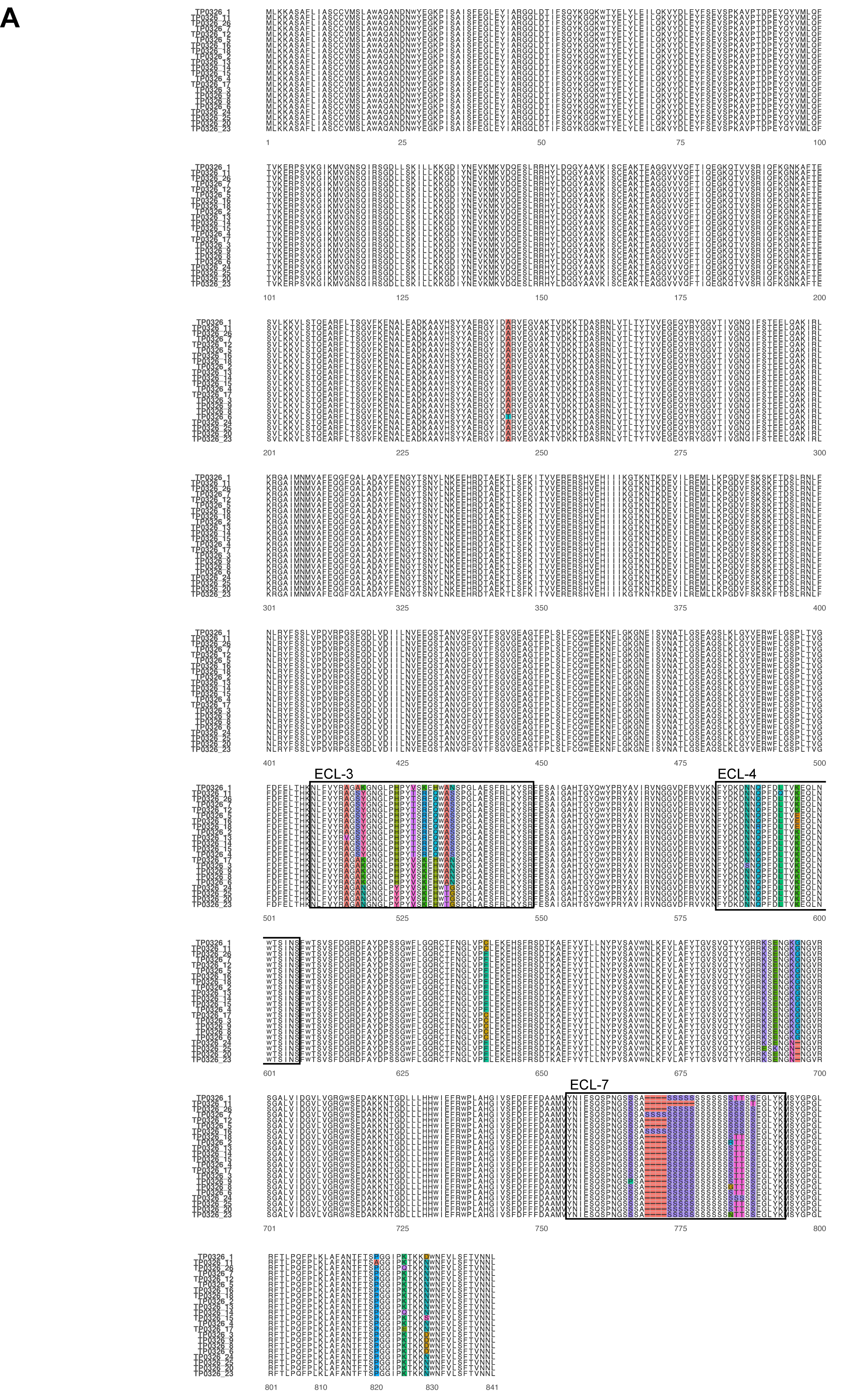

### Figure S2BCD

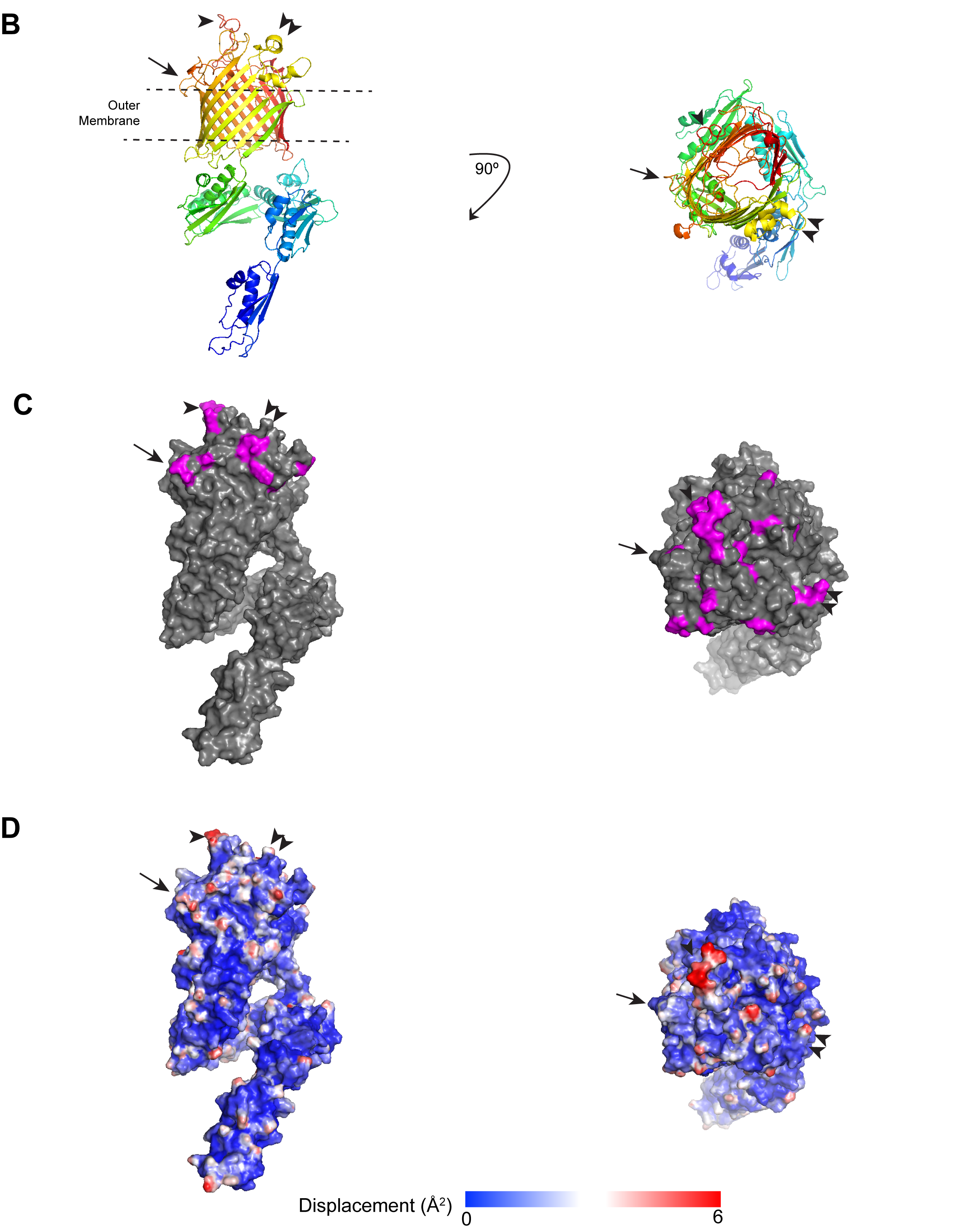

### Figure S3A

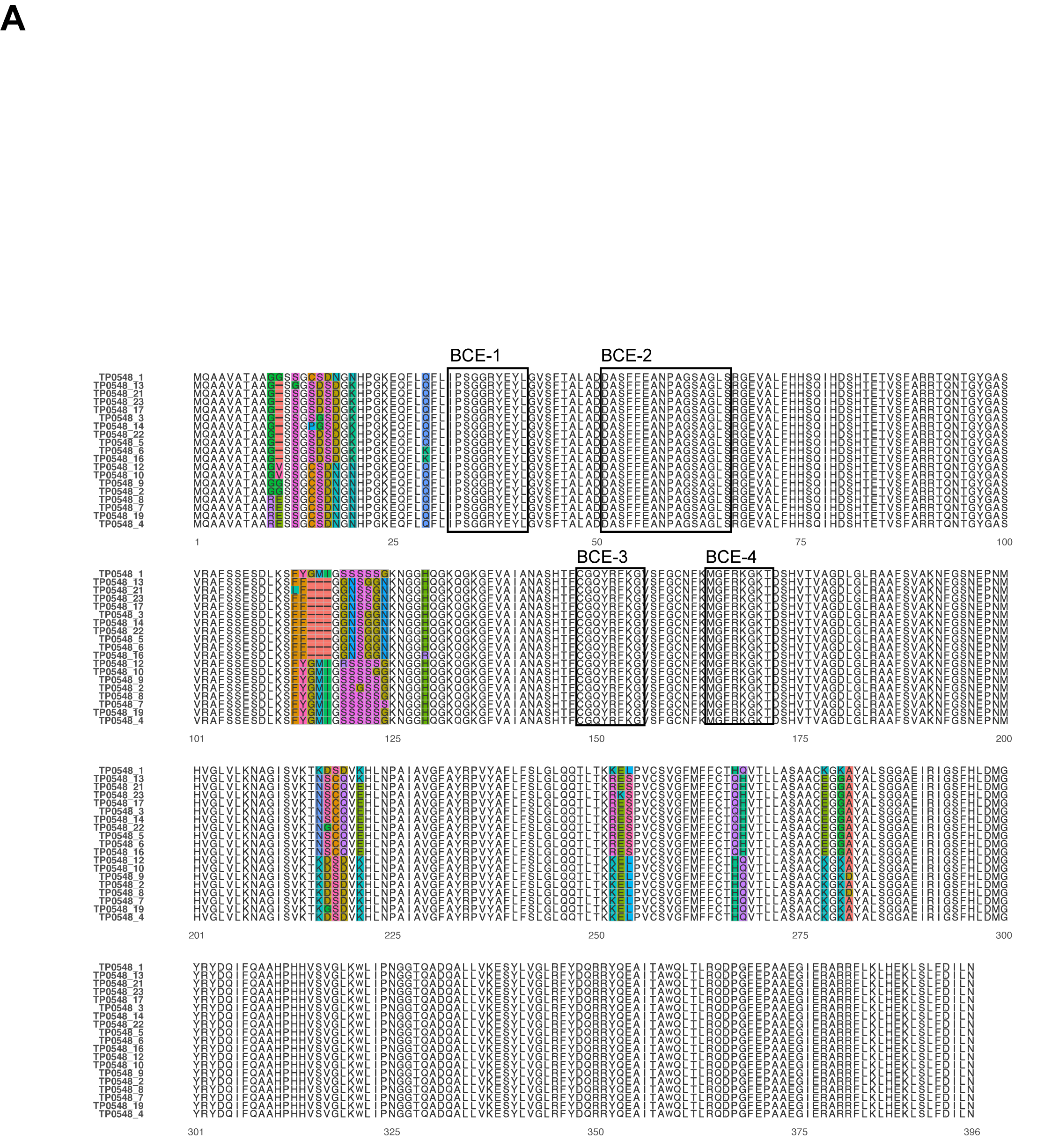

### Figure S3BCD

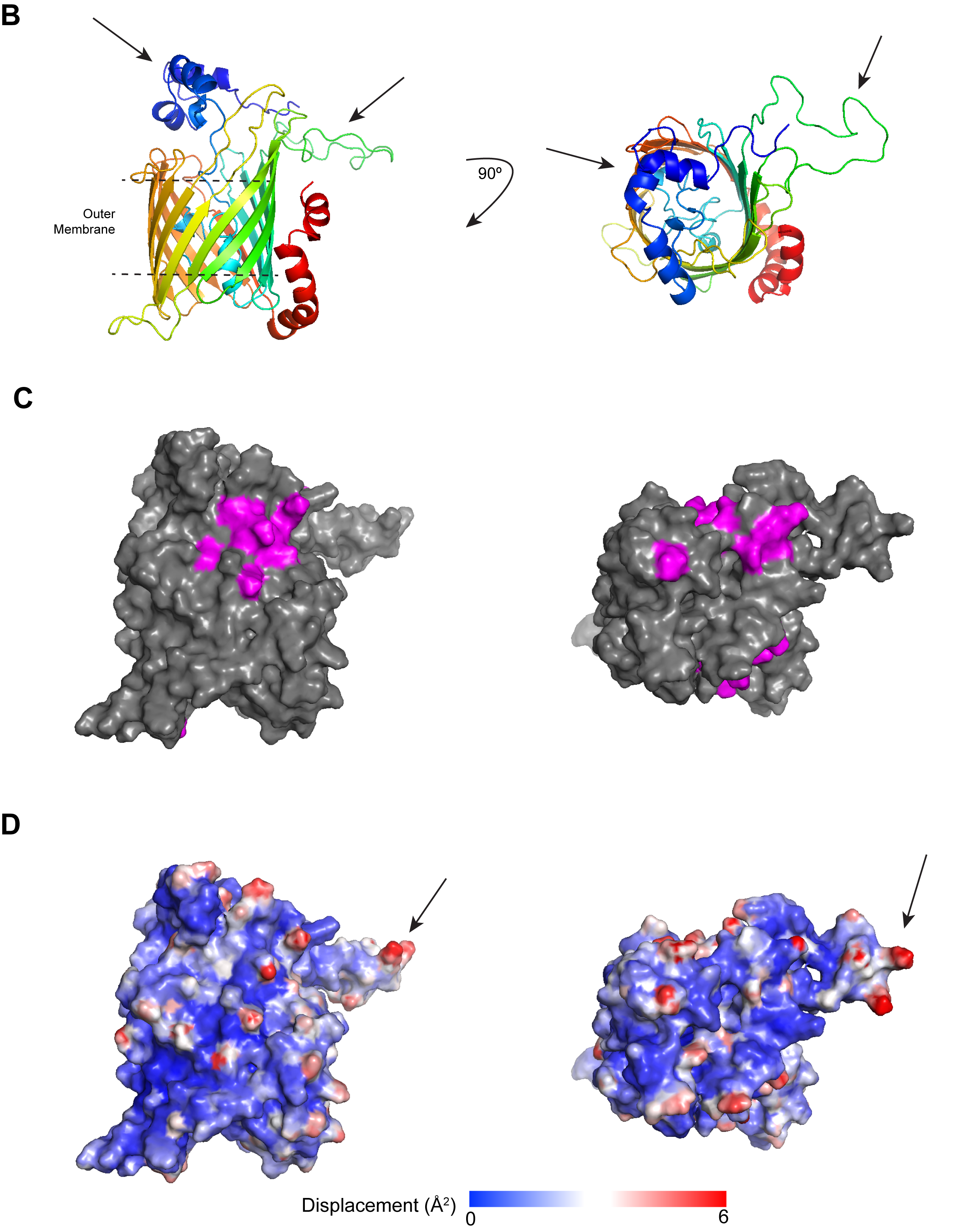

### Figure S4A

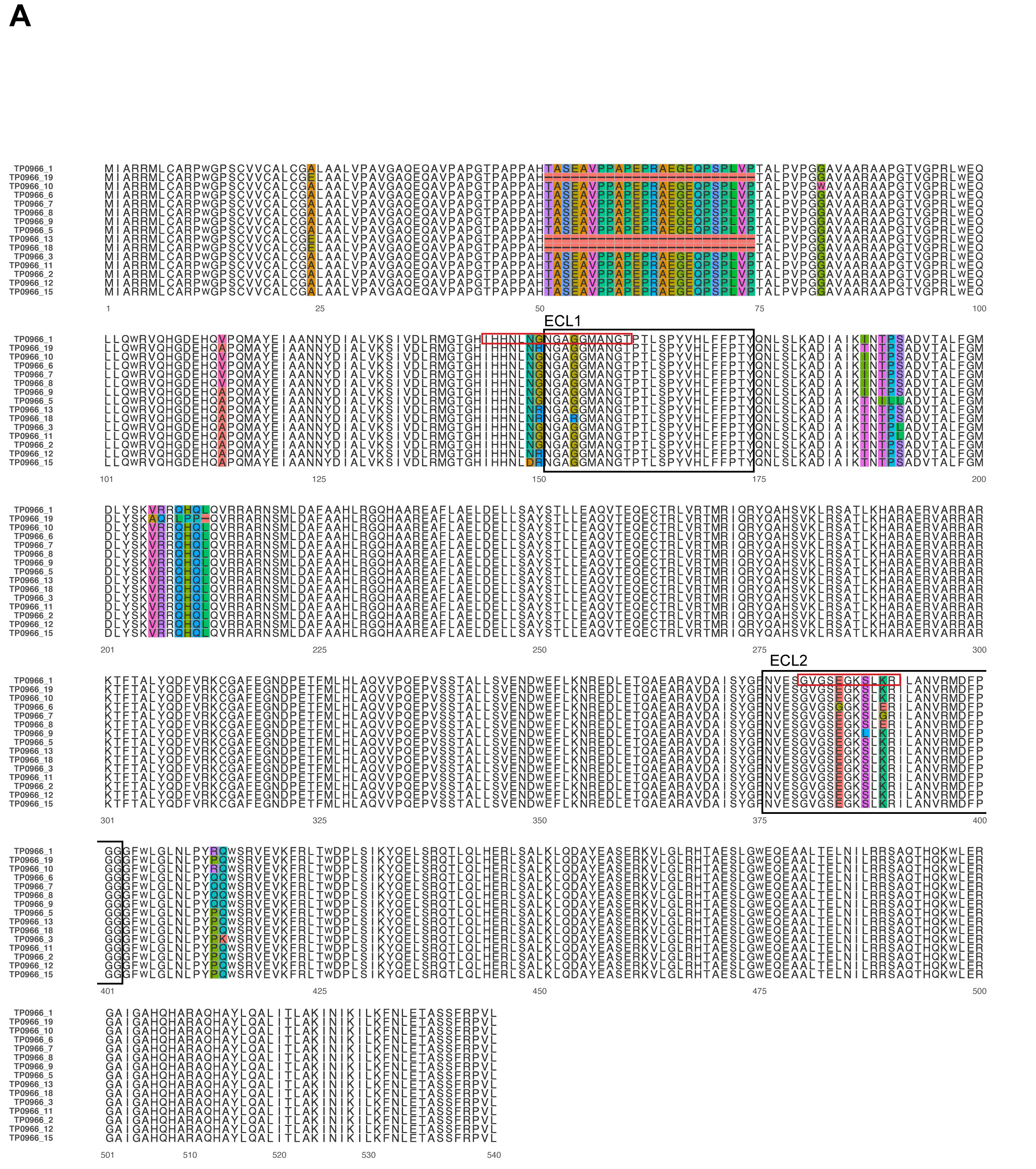

### Figure S4BCD

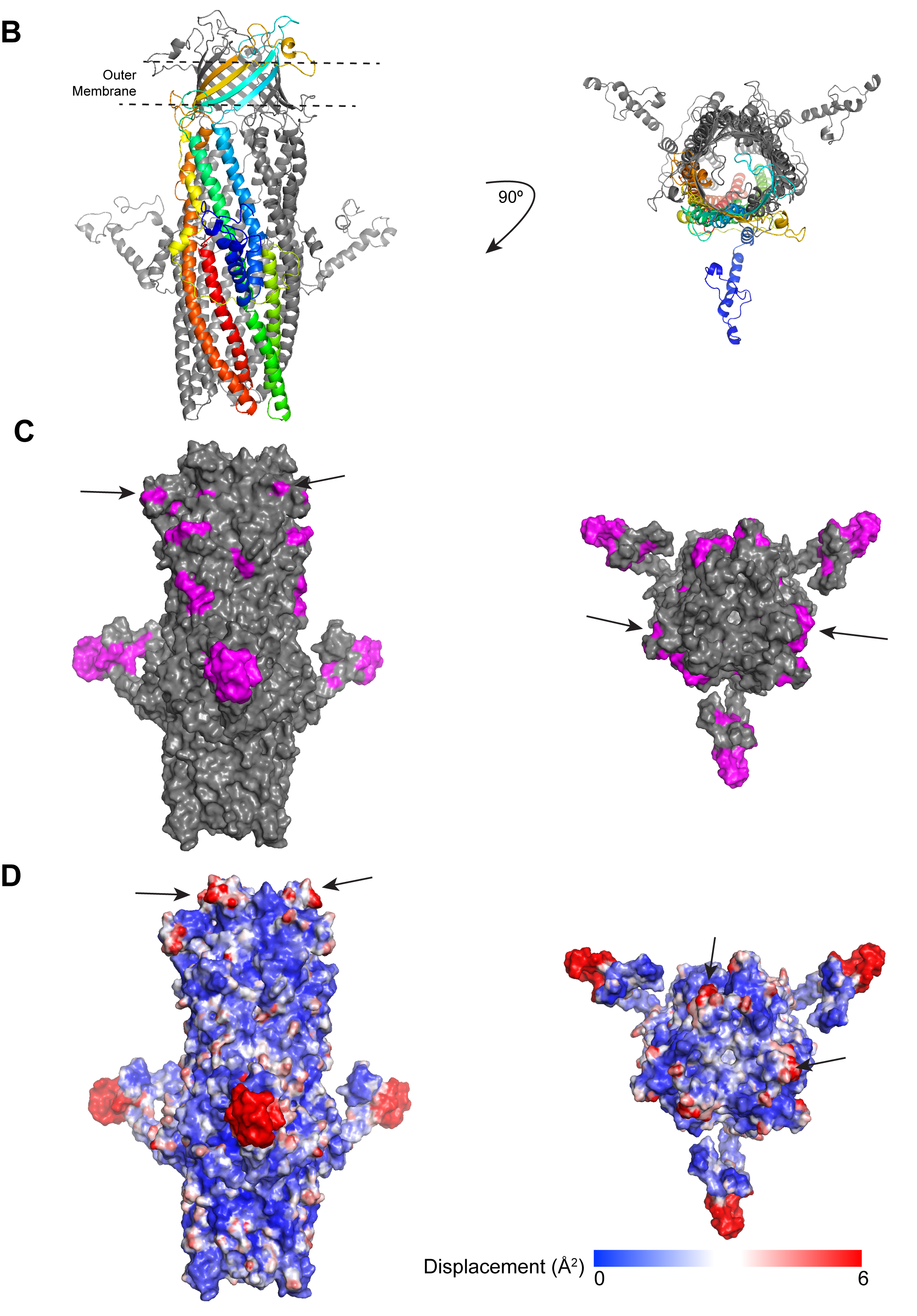

### Figure S5A

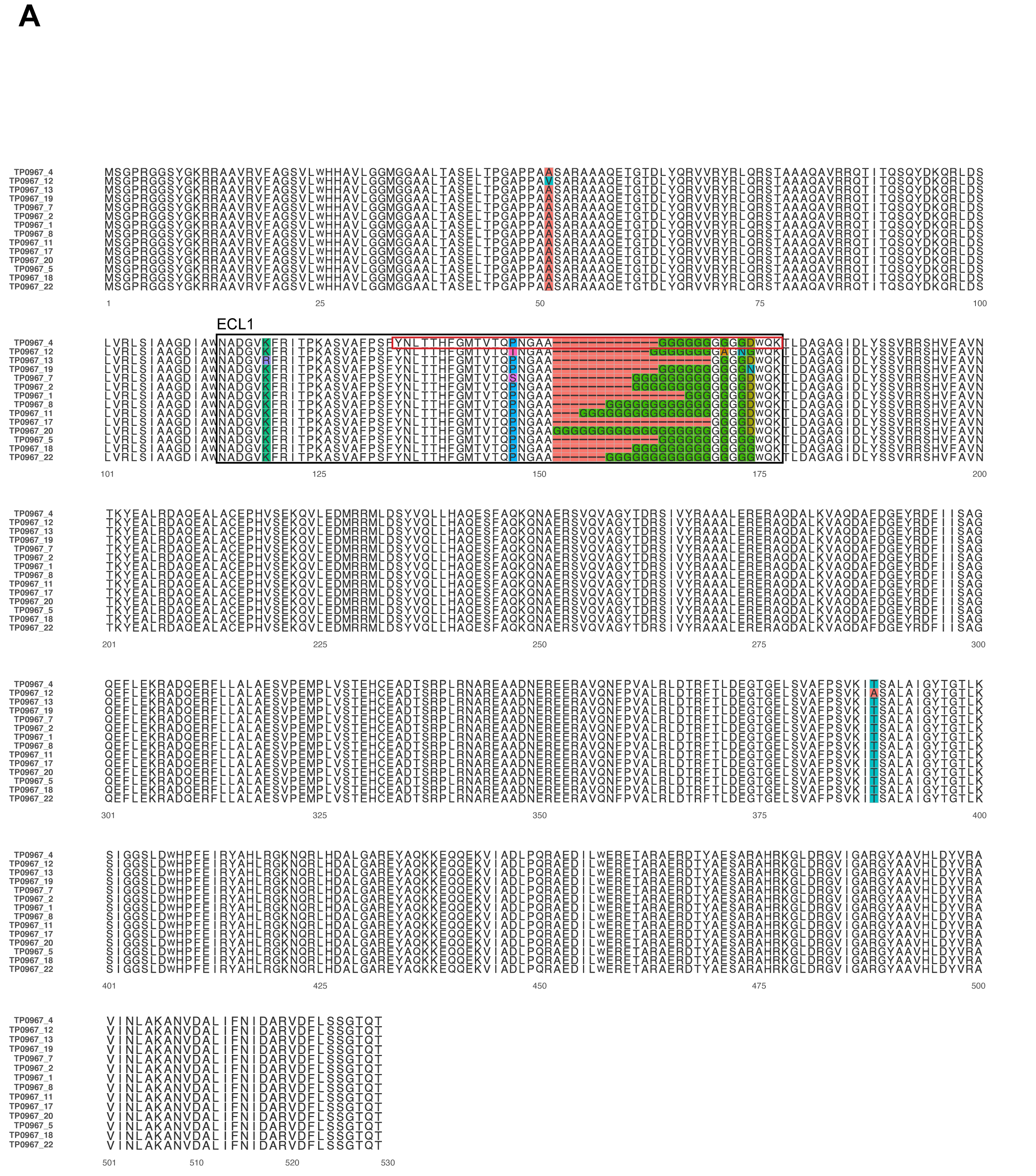

### Figure S5BCD

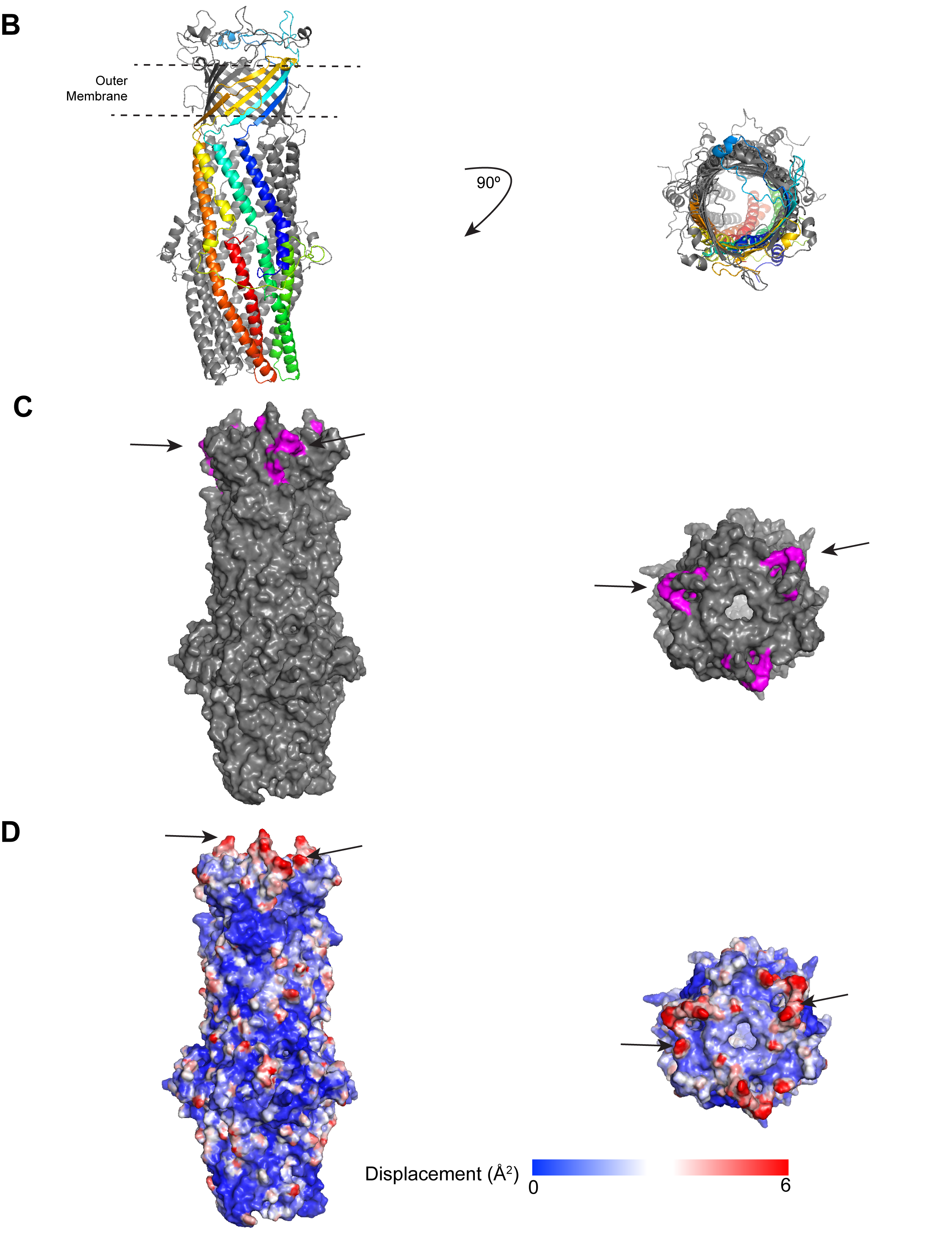

### Figure S6A

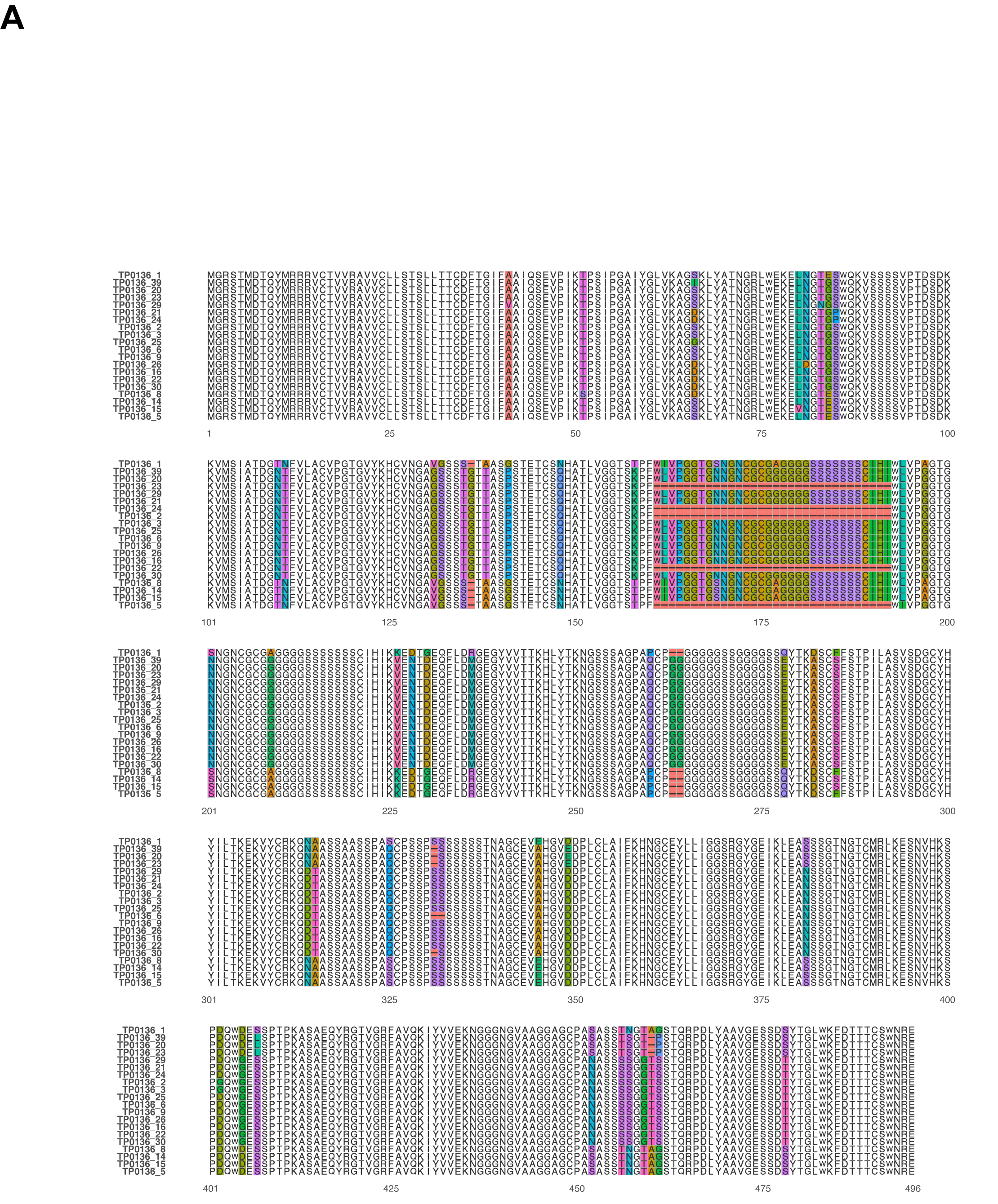

### Figure S6BCD

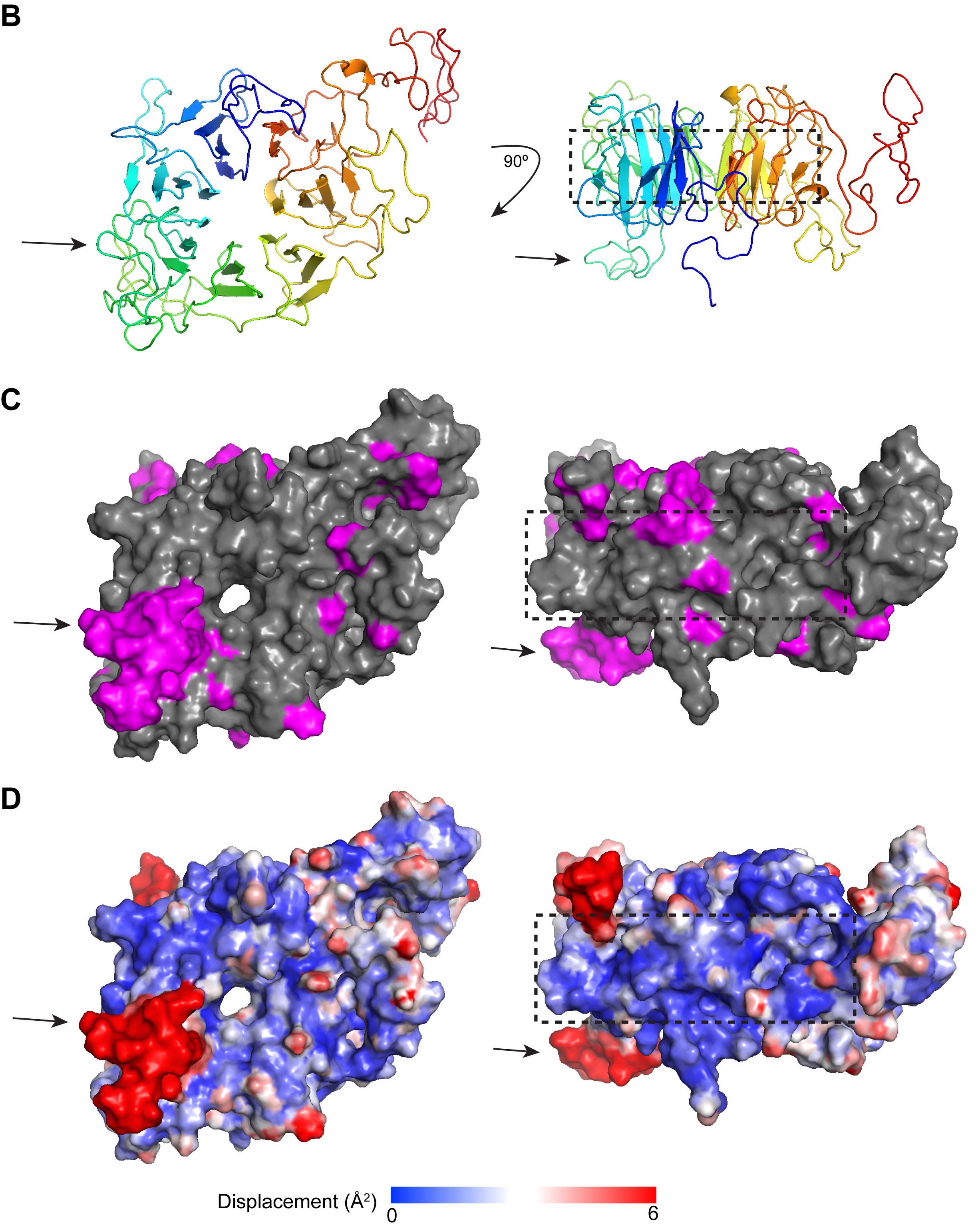
